# Determinants of transcription initiation efficiency and start site selection by RNA polymerase I

**DOI:** 10.1101/2024.10.30.621142

**Authors:** Olena Parilova, Piia Bartos, Anssi M. Malinen

**Affiliations:** Department of Life Technologies, University of Turku, Turku, Finland; School of Pharmacy, University of Eastern Finland, Finland

## Abstract

RNA polymerase I (Pol I) synthesizes pre-ribosomal RNA, which is essential for ribosome biogenesis. Increased Pol I activity supports rapid cell growth, a key characteristic of cancer cells. Consequently, drugs targeting Pol I in cancer are being actively investigated. The first step in pre-rRNA synthesis involves the assembly of the Pol I transcription initiation complex on the rRNA gene promoter, yet the quantitative and dynamic parameters of this process remain inadequately defined. We combined biochemical, biophysical, and molecular dynamics simulation analyses to enhance molecular models illustrating how Pol I apparatus isolated *Saccharomyces cerevisiae* recognizes the promoter and the transcription start site (TSS). Time-resolved binding data shows that CF relies on a two-step mechanism, consisting of the binding and isomerization steps, to recognize its binding site on the promoter. The next step, CF-mediated recruitment of the Pol I– Rrn3 complex to the promoter, was inefficient, leading to the rapid dissociation of Pol I. The selection of the TSS by Pol I is influenced by the distance to CF and the properties of downstream DNA, such as bendability. The dynamics revealed in the Pol I transcription apparatus establish a framework for comparing the functions and mechanisms of Pol I activators and inhibitors.

## INTRODUCTION

Cell growth and homeostasis rely on efficient protein synthesis, which is heavily influenced by the number of functional ribosomes ^1,2^. A critical checkpoint in ribosome production is the transcription of ribosomal RNA genes (rDNA), responsible for over 60% of total RNA synthesis in growing cells^3^. Transcription is key process in regulating ribosome synthesis, responding to various environmental signals related to growth control. Transcription of the multicopy rDNA gene locus is carried out by RNA polymerase I (Pol I) and RNA polymerase III. An oversupply of ribosomes is linked to accelerated cellular proliferation, while a shortage can induce cell cycle arrest, senescence, or apoptosis ^4^. Dysregulation of Pol I transcription and impaired ribosome biogenesis are also implicated in numerous diseases, including ribosomopathy syndromes, developmental disorders and many cancer types ^1,4–6^. Cancer cells hijack transcription regulatory points to permanently activate rRNA synthesis ^1,3^. Further, genes encoding core components of the Pol I transcription machinery are frequently upregulated in some cancers, e.g., breast invasive carcinoma, esophageal carcinoma, lung adenocarcinoma and testicular cancer ^7^.

In *Saccharomyces cerevisiae* (yeast), the Pol I transcription system is homologous to that in mammals, making yeast a valuable model for studying fundamental aspects of eukaryotic transcription by Pol I. Yeast Pol I transcribes rDNA to 35S precursor rRNA, which is processed to 18S, 5.8S, and 25S rRNA components of the ribosome. The mature ribosome additionally contains 5S rRNA, synthesized by RNA polymerase III. While the structural basis of the Pol I machinery, including the initiation complex, has been recently extensively studied using cryo-electron microscopy (cryo-EM) ^8–12^, the functional aspects require further investigation. Detailed studies of Pol I function can decipher the exact molecular mechanisms that control rDNA transcription and identify novel druggable targets in the Pol I apparatus and its regulators.

Transcription of rDNA begins with the formation of the closed complex (CC) on the promoter DNA, characterized by the recruitment of Pol I to the promoter region proximal to the transcription start site (TSS) through transcription factors UAF, TBP, CF, Rrn3, and NET1. In *S. cerevisiae*, the multifunctional Net1 protein was reported to stimulate Pol I transcription that can be attributed to the C-terminal region (CTR) of Net1, which was required for normal cell growth and Pol I recruitment to rRNA genes *in vivo* and sufficient to promote Pol I transcription *in vitro* ^13^. Net1 directly binds Pol I and stimulates rRNA synthesis both *in vitro* and *in vivo* ^14^. Additionally, Net1 modulates nucleolar structure by regulating rDNA morphology ^14^. UAF is multi-subunit complex that regulate preinitiation complex formation by binding to the upstream activating sequence at the rDNA promoter and by stimulating recruitment of additional Pol I factors to the more downstream positions on the promoter. Yeast UAF contains histones H3 and H4 and four nonhistone protein subunits. TBP was originally identified as a protein component of TFIID that binds directly to TATA box present in many Pol II promoters. Later, TBP was found to be required not only for the initiation of transcription by Pol II but also for the initiation of transcription by Pol I and Pol III ^15^. The requirement of TBP for rDNA transcription by Pol I was demonstrated for mammalian ^16,17^, yeast ^18,19^ and amoeba (*Acanthamoeba castellanii*) ^20–22^ transcription systems. However, in contrast to the role of TBP in transcription of TATA-containing promoters by Pol II, the function and mechanism of TBP in TATA-less promoters, including the rDNA promoter, were less defined for a long time. *In vitro* yeast rDNA transcription and binding experiments, and recent cryo-EM structures of preinitiation complex uncovered the role of TBP. TBP can form a complex with CF and simultaneously also interact, even more strongly, with UAF ^23^. Consistent with *in vitro* observations, CF also appears to be indispensable for rDNA transcription *in vivo*, and the genes encoding CF subunits, specifically *RRN6*, *RRN7* and *RRN11*, are all essential for cellular growth ^24,25^. Early biochemical studies reported that purified CF does not form a stable complex with promoter DNA by itself but becomes part of a stable preinitiation complex in the presence of other transcription factors ^24^. Notably, CF subunit Rrn6 has strong interaction interface with TBP ^10,18,26^ while Rrn7 subunit interacts with both UAF and TBP ^18^. Rrn3 is an essential protein required for rDNA transcription by Pol I. Rrn3 interacts with Pol I in the absence of DNA template and stimulates recruitment of Pol I to the promoter by providing Pol I additional physical connection to CF ^27^. Contrary to CF and Rrn3, the essential nuclear factors for rRNA synthesis *in vitro* and *in vivo*, UAF, TBP and Net1 are stimulatory but not essential. *In vitro* the addition of TBP without UAF fails to stimulate basal transcription ^28^. Thus, TBP mediates activated transcription from the rDNA promoter in association with UAF ^19^. These differences define two groups of transcription factors reflecting their core (basal) or stimulatory (accessory) function in rDNA transcription. Indeed, the basal yeast protein machinery for promoter-specific transcription initiation consists of only Pol1, CF and Rrn3. CF binds a core promoter element upstream of the TSS and then recruits the Pol I·Rrn3 complex, enabling a basal rate of transcription initiation ^29,30^. DNA is loaded into the expanded active center cleft of the polymerase. DNA unwinding and transcription bubble formation between the protrusion and clamp domains of Pol1 enables cleft contraction, resulting in an active Pol I and promoter DNA conformation known as an open complex (OC). The beginning of RNA synthesis marks the transition into an initially transcribing complex (ITC), which undergoes further structural rearrangements, including the breakage of Pol I interaction with CF and Rrn3, as the final transformation to an elongation complex (EC), characterize by productive RNA synthesis downstream from the promoter. The suggested mechanism of Pol I specificity to rDNA promoter relies on a distinct ‘‘bendability’’ and ‘‘meltability’’ of the promoter sequence that enables contacts between initiation factors, DNA and polymerase ^9,11^.

Here, we combined biochemical, biophysical and molecular dynamic simulation analyses to study the basal yeast Pol1 transcription initiation apparatus recognizes the rDNA promoter and determines the TSS. Our time-resolved binding data shows that CF utilizes a two-phase mechanism, consisting of separate binding and isomerization steps, to recognize its binding site on the rDNA promoter. The stability of CF·promoter complex largely depends on the sequence-specific interaction at the upstream edge of the CF binding site. Promoter-bound CF then recruits Pol1·Rrn3 holoenzyme to the promoter, however, the formed preinitiation complex is not stable, leading to continuous Pol1 binding and dissociation dynamics. Pol1 lacked the capability to recognize specific bases near the TSS and instead appears to select the TSS based on the interaction distance with the promoter-bound CF and the physical properties of the DNA, i.e., “bendability” of the DNA segment near the upstream edge of the forming transcription bubble.

## MATERIALS AND METHODS

### Protein production and purification

Pol I purification followed a previously published protocol with some modifications ^31^. Pol I was extracted from *Saccharomyces cerevisiae* SC1613 strain (constructed by Cellzome AG, Germany), which has the chromosomal gene for AC40 subunit modified to encode a C-terminal tandem affinity purification (TAP) tag. The TAP tag consists of a calmodulin binding domain, a TEV protease recognition site and a protein A domain ^32^. The genotype of *S. cerevisiae* SC1613 is MAT a; ade2; arg4; leu2-3, 112; trp1-289; ura3-52; YPR110c::TAP-K.I.URA3. To produce yeast biomass, an inoculum was started from a glycerol prep and grown in a shaker cultivator (set to 200 rpm and 30°C) in Erlenmayer flask containing 50 mL of synthetic dropout media made of yeast nitrogen base, without aminoacids, without ammonium sulphate (Alfa Aesar, cat. no H26271), lacking uracil but supplemented with auxotrophic amino acids, ammonium suphate and 2% glucose. When the cell density reached absorbance value 4.0–4.5 at 600 nm (A_600_), the pre-culture was used to seed 22 L of YPD medium in Bioengineering NLF22 bioreactor (Bioengineering AG; vessel size 30 L) to a starting cell density (A_600_ ∼0.05). Yeast cells were propagated at 30℃, with the stirring speed set to 250 rpm and aeration level to 18–22 L/min, until A_600_ reached 2.5–3.2. Cells were then harvested by centrifugation, washed once with ice-cold water, snap frozen in liquid nitrogen and stored at - 80℃ until use.

To purify Pol I, 80–145 g of frozen yeast cells were resuspended in binding buffer A [250 mM Tris-HCl, pH 8.0, 40% (v/v) glycerol, 250 mM (NH_4_)_2_SO_4_, 12 mM β-mercaptoethanol, 10 mM MgCl_2_, 1 mM EDTA, and 10 μM ZnCl_2_] supplemented with protease inhibitors (Pierce™ Protease Inhibitor Mini Tablets, EDTA-free; ThermoFisher Scientific; cat.no. A32955). The cells were disrupted using 0.5 mm diameter glass beads and BeadBeater (BioSpec). After the removal of glass beads by pouring the sample through a miracloth filter (Millipore), the lysate was treated with DNase I (5 μg/ml) and clarified by centrifugation at 64000 g for 1 h at 4 ℃ using JA-25.5 rotor. The supernatant was aliquoted to glass Econo-Columns (diameter 2.5 cm, height 20 cm, BioRad) each containing 3 mL of Heparin Sepharose resin (GE Healthcare), followed by incubation on a rocking shaker for 2.5 h at 6℃. The resin was washed with 5 bed volumes (CV) of buffer B [50 mM Tris-HCl, pH 8.0, 20% (v/v) glycerol, 250 mM (NH_4_)_2_SO_4_, 1 mM MgCl_2_, 1mM β-mercaptoethanol, 0.5 mM EDTA, 10 μM ZnCl_2_] to remove proteins loosely bound to heparin resin. Next, Pol I containing fraction was eluted with about 3 CV of buffer C [buffer B supplemented with total 1M (NH_4_)_2_SO_4_]. Pol I containing eluate was diluted to 500 mM (NH_4_)_2_SO_4_ using buffer D [buffer B with 0 mM (NH_4_)_2_SO_4_, 0,1 % (v/v) Tween-20] and incubated with at least 1.2 mL of pre-equilibrated IgG Sepharose (GE Healthcare) for 6 h at 6℃ in a rotating shaker. IgG resin was then washed with buffer E [50 mM Tris-HCl, pH 8.0, 20% (v/v) glycerol, 225 mM (NH_4_)_2_SO_4_, 1 mM MgCl_2_, 0.5 mM EDTA, 2 mM β-mercaptoethanol, 10 μM ZnCl2, and 0.05% (v/v) Tween-20]. Pol I complex was then recovered from the resin by cleaving the TAP-tag with TEV protease in 1 CV of buffer E [≥150 µg/ml TEV, 2 h at 16°C, shaking 1000 rpm in a thermo-shaker (Grant)]. After cleavage, eluate was collected by spinning down the resin (500g, 3 min, 4°C). The harvested resin was washed with 10 CV of recovering buffer F [50 mM Tris-HCl, pH 8, 60 mM ammonium sulfate, 0.5 mM EDTA, 1 mM MgCl2, 10 μM ZnCl2, 2 mM b-mercaptoethanol, 0.05% (v/v) Tween-20] and next combined with eluate. The eluted sample containing Pol I and Pol III protein fractions was loaded into a 1 mL Resource Q anion exchange column (GE Healthcare), which was then eluted, using Äkta Purifier (GE Healthcare) and flow rate 1 mL/min, with a gradient of 0.06–1 M (NH_4_)_2_SO_4_ in 30 mL of buffer G [40 mM Tris-HCl, pH 8.0, 5 mM DTT, 1 mM MgCl_2_, 0.5 mM EDTA, and 10 μM ZnCl_2_]. Purified Pol I was collected at ∼280 mM (NH_4_)_2_SO_4_ and concentrated with a centrifugal filtration unit (Amicon Ultra-4, 30 kDa cutoff, Millipore) to 2–7 μM final concentration and supplemented with 15% (v/v) glycerol. Protein fractions and final Pol I preparations were quality controlled by SDS-PAGE and Western analysis using Coomassie staining and anti-TAP-tag antibody (TAP Tag Monoclonal Antibody DyLight™ 800 4X PEG, clone 22F12-1E3, cat.no. 15342907, Fisher Scientific), respectively.

The production and purification protocol of Rrn3 was modified from a previous report ^30^. We cloned the gene of Rrn3 by PCR from the genomic DNA of *Saccharomyces cerevisiae* INVSc1 (ThermoFisher Scientific) and ligated it into pET-15 to prepare the expression plasmid pAM037, which carries ampicillin resistance marker (**Table S1**). Rrn3 was engineered to contain C-terminal TEV cleavable 10xHis tag. pAM037 was transformed into *Escherichia coli* B95.ΔA strain ^33^. Cells were initially grown in a shaker incubator (set to 170 rpm) at 37℃ in six 2 L Erlenmeyer flasks each containing 0.8 L of LB medium supplemented with 50 mg/L ampicillin. When A_600_ reached 1.0, the culture was cooled to 20°C, supplemented with 0.2 mM IPTG to induce Rrn3 production and further incubated overnight. Cell suspension was harvested by centrifugation (4500 g, 20 min, 4℃), flash-frozen in liquid nitrogen and stored at -80℃ until use. Cell mass was thawed in buffer H [50 mM Hepes, pH 7.8, 500 mM NaCl, 20 mM imidazole, 5 mM MgCl_2_, 1 mM β-mercaptoethanol, 0.1 mM EDTA, 10% (v/v) glycerol, 0.2% Tween-20], supplemented with 0.125 mg/ml lysozyme and protease inhibitors (Pierce™ Protease Inhibitor Mini Tablets, EDTA-free; ThermoFisher Scientific; cat.no. A32955). Cells were disrupted by sonication using Soniprep 150 (MSE) in 30 s on/off duty cycle and ice-water bath to ensure efficient cooling. Cell lysate was cleared by centrifugation (75 600 g, 1 h, +4°C) and supernatant was loaded into a 5 mL Nickel sepharose column (HisTrap HP, GE Healthcare). The loaded column was first washed, using Äkta purifier FPLC system (GE Healthcare), with 20 mM imidazole in buffer J [50 mM Hepes-KOH, pH 7.8, 500 mM NaCl, 5 mM MgCl_2_, 0.5 mM β-mercaptoethanol, and 10% (v/v) glycerol] followed by a more stringent wash with 40 mM imidazole in buffer J. Protein was then eluted using a steep imidazole gradient (40–500 mM imidazole in 25 mL of buffer J, flow rate 1 mL/min). Eluate was diluted in 1:3 ratio with buffer K [20 mM HEPES, pH 7.8, 5 mM MgCl_2_, 1 mM β-mercaptoethanol, 0.1 mM EDTA, and 5 % (v/v) glycerol] and applied into a 1 mL Resource Q anion exchange column. Rrn3 was eluted with a gradient of 0.225–1.5 M NaCl in buffer K using Äkta FPLC (GE Healthcare). Rrn3 eluted from the column at ∼0.45 M NaCl. Final protein preparation was concentrated using a centrifugal filter unit (Amicon Ultra-4, 3 kDa cutoff, Millipore) at 3500 g, 4℃, and exchanged to buffer L [20 mM HEPES, pH 7.8, 150 mM NaCl, 1 mM MgCl_2_, 0.1 mM EDTA, 0.1 mM DTT, and 50% (v/v) glycerol] using Slide-A-lyzer Mini dialysis device (10 K MWCO, 0.5 mL, ThermoFisher Scientific). Fractions and final Rrn3 preparations were quality controlled by SDS-PAGE and Western analysis using Coomassie staining and monoclonal anti-His-tag antibody ( 6X His Tag Antibody Dylight™ 800 Conjugated, clone 33D10.D2, cat.no. 200-345-382, Rockland Immunochemicals), respectively.

Recombinant core factor (CF) protein was produced using an approach similar to previous reports ^29,34^. We cloned the genes of CF subunits from the genomic DNA of *S. cerevisiae* INVSc1 (ThermoFisher Scientific). The genes of Rrn7 and Rrn11 were ligated into pET-36b plasmid (Novagen) to prepare expression plasmid pAM033, which carries kanamycin resistance marker. The gene of the third CF subunit, Rrn6, was cloned into pET-15 with C-terminal TEV cleavable 10xHis tag to prepare expression plasmid pAM031, which carries ampicillin resistance marker. pAM031 and pAM033 (**Table S1**) were co-transformed into *E. coli* XJb (DE3) (Zymo Research) or T7 express (New England Biolabs) cells for the production of CF. The XJb cells were also transformed with pRIL [isolated from *E. coli* BL21(DE3)-RIL (Stratagene)], a plasmid that supplies for the production strain rare transfer-RNA’s for AGA/AGG, AUA and CUA codons, and carries chloramphenicol resistance marker ^35^. *E.coli* were grown in 2 L Erlenmeyer flasks (6–8 pieces) each containing 0.8 L of LB medium supplemented with 100 mg/L ampicillin 30 mg/mL kanamycin and 30 mg/mL chloramphenicol; the shaker incubator was set at 37°C and 170 rpm. When A_600_ reached 0.8, protein production was induced with 0.2 mM IPTG. After overnight induction at 20°C, cells were harvested by centrifugation (4500 g, 20 min, 4℃), flash-frozen in liquid nitrogen and stored at -80℃ until use. Cell mass was resuspended in buffer F supplemented 0.125 mg/ml lysozyme and protease inhibitors (Pierce™ Protease Inhibitor Mini Tablets, EDTA-free; ThermoFisher Scientific; cat.no. A32955). Cells were disrupted and CF captured from the lysate by metal-affinity chromatography as described above for Rrn3. The CF containing eluate from Nickel sepharose column was diluted in 1:2 ratio with buffer K to decrease NaCl to 170 mM concentration. Benzonase nuclease was added to 50 U/ml, followed by 30 min incubation under moderate rotation at 6°C. The sample was then loaded into a 5 mL HiTrap Heparin HP column (GE Healthcare) at flow rate 1.5 mL/min. Elution was performed with a gradient of 0.225–1.5 M NaCl in buffer K using Äkta FPLC (gradient length 30 mL, flow rate 1 mL/min, GE Healthcare). CF eluted at ∼825 mM NaCl. CF containing fractions were combined and concentrated at 4℃ using Amicon Ultra-4 centrifugal filter unit (30 kDa cutoff, Millipore). The final CF preparation was exchanged to buffer M [20 mM HEPES, pH 7.8, 1 M NaCl, 1 mM MgCl_2_, 0.1 mM EDTA, 50% (v/v) glycerol, 0.1 mM DTT] using Slide-A-lyzer Mini dialysis device (10 K MWCO, 0.5 ml, ThermoFisher Scientific). Purification fractions and final CF preparations were quality controlled by SDS-PAGE and Western analysis using Coomassie staining and monoclonal anti-His-tag antibody ( 6X His Tag Antibody Dylight™ 800 Conjugated, clone 33D10.D2, cat.no. 200-345-382, Rockland Immunochemicals), respectively.

### Size exclusion chromatography of protein preparations

Transcription factor preparations used in biophysical assays (mass photometry and protein induced fluorescence enhancement) were further polished using size-exclusion chromatography. Prior to the gel filtration step the His-tag in CF was cleaved off by supplementing concentrated CF sample (vol. ≤500 µl) with TEV protease and incubating the sample either overnight at 4°C or 2 h at 16°C in a thermo-shaker (Grant Instruments) set to 1000 rpm. A Superose 6 Increase 10/300 GL gel filtration column (Cytiva) was pre-equilibrated with GF-CF buffer [20 mM HEPES, pH 7.8, 300 mM NaCl, 1 mM MgCl_2_, 0.1 mM EDTA, and 0.1mM DTT] using Äkta purifier FPLC (GE Healthcare). TEV-treated CF sample (∼500 µL) was injected into the column and run at 0.4 mL/min at 10 °C using GF-CF buffer. Resolved proteins were collected in 500 µL fractions and analyzed by NanoDrop spectrophotometer (ThermoFisher Scientific) and SDS-PAGE. Size-exclusion chromatography of Rrn3 was carried out similarly to CF with the exception that GF-R buffer [20 mM HEPES, pH 7.8, 150 mM NaCl, 1 mM MgCl_2_, 0.1 mM EDTA, 0.1 mM DTT] was used. Final preparations of CF and Rrn3 were concentrated using centrifugal filters (Amicon Ultra-4, Millipore), stabilized with the addition of glycerol to final concentration 20% (v/v), aliquoted, and flash frozen in liquid nitrogen for storage at -80°C.

### DNA template preparation

Oligonucleotides encoding wild-type or mutant *S. cerevisiae* rDNA promoters were purchased from Merck or Eurofins Genomics. To prepare double stranded promoter scaffolds, the equimolar amounts of template and non-template strands (5–10 µM) were mixed in 0.1% (v/v) DEPC treated buffer [10 mM HEPES-KOH, pH 7.5, 50 mM NaCl, and 100 µM EDTA] and annealed by using a PCR machine at 94℃, 3 min to gradually cool the mixture from 94℃ to 4℃. The ready-to-use scaffolds were stored at -20℃.

To prepare circular supercoiled transcription templates, rDNA promoter fragment from position -200 to +150 [relative to the TSS at +1] or promoter-free fragment from ATG13 gene (length 371 bp) was cloned by PCR from the genomic DNA of *S. cerevisiae* INVSc1 (ThermoFisher Scientific) and ligated between SacI and HindIII restriction sites in pUC18 plasmid (**Table S1**). Plasmids, pAM036 (contains rDNA promoter) and pAM041 (ATG3 fragment) were maintained and amplified using *E. coli* XL1-blue cells. The plasmids were isolated using GeneJET Plasmid Miniprep Kit (ThermoFisher Scientific) according to the manufacturer’s protocol and stored at -20°C. The plasmids were verified by Sanger sequencing the insert and the insert–plasmid junctions.

The plasmid concentrations were determined using NanoDrop spectrophotometer, and the plasmids were stored at -20°C. The linear scaffolds of promoter (−200/+150) and non-promoter DNA (ATG3) were produced by PCR amplification of the target fragment from pAM036 and pAM041 vectors (**Table S2**). The DNA sequences of all transcription template constructs are given in **Table S3**.

### In vitro transcription

*In vitro* transcription reactions were performed as previously described with modifications ^36,37^. Plasmids (circular negatively supercoiled) and linear templates containing either native, truncated native, base-substituted, pre-melted or forked core element of rDNA promoter were examined. Basal preinitiation complex (PIC) was reconstituted by first mixing 2.4 pmol of synthetic DNA template with 15 pmol of CF (medium-pure preparation) and incubating the mixture in transcription TIB10 buffer (TIB [100 mM HEPES/50 mM KOH, pH 7.5, 100 mM K-glutamate, 0.025 mM ZnCl2, 5% (v/v) glycerol, 1 mM EDTA, 0.2 mM TCEP, DEPC treated] supplemented with 10 mM Mg acetate) for 15 min at 22 °C. Then, a pre-incubated holoenzyme complex (6 pmol Pol I and 12 pmol Rrn3) in TIB2 was added in ratio 1:1 and the incubation was continued for another 15 min at 30℃. The transcription reaction was initiated by the addition of equal volume of 1 mM each NTP (ATP/CTP/GTP/UTP) in TIB10 to PIC. The final volume of transcription reaction mixture was 10 µL and contained 1 pmole template DNA, 2.5 pmole Pol I, 5 pmole Rrn3, 6.25 pmole CF, and 5 nmole (0.5 mM) NTPs in TIB8; the reaction tube was incubated in a thermo-mixer (Grant) at 30°C for 20 min. Control transcription reactions without NTP addition were prepared in parallel. Transcription reaction was terminated by the addition of 1 µl 105 mM EDTA (to ∼9.5 mM final concentration) and immediate heating of the samples at 75°C for 5 min and transferred on ice.

The quantity and length from 5’ end (i.e., the TSS of RNA products) were determined using primer extension. The inactivated transcription reactions were supplemented with 1.25–6 pmol of fluorophore-labelled DNA primer (OP003, OP021 or primarily OP018, see **Table S2**) and with Mg^2+^ (1 µl of 150 mM MgCl_2_ stock) to gain 5 mM of unchelated Mg^2+^ concentration. The primer was annealed to the target site of the RNA transcripts by heating the samples at 95°C for 80 s followed by cooling on ice for 15 min. Reverse transcription reaction was initiated by the addition of 12 µl of mixture containing 100 U SuperScript II Reverse Transcriptase (Invitrogen), 0.5 mM dNTPs, 5mM DTT, 10 U Murine RNase inhibitor (New England Biolabs), and 50 µg/ml Actinomycin D in 1x SuperScript buffer. The reactions (total vol. ∼25 µl) were incubated at 47°C for 1 h, and then terminated by the addition of formamide gel loading buffer (92.5% formamide, 0.039 M LiOH, 0.013 M EDTA, 0.23% OrangeG) in 1:1 ratio followed by heating at 95℃ for 3 min. The single stranded DNA products of reverse transcription reaction were separated on 12% denaturing polyacrylamide gels and visualized with an Odyssey Infrared Imager (Li-Cor Biosciences) at 700 nm channel. The TSS of transcription reaction products was determined by comparing their electrophoretic mobility to that of the manual Sanger sequencing products of wild-type rDNA promoter run on the same gel. Sequencing reactions were done using Cy5.5-labeled OP018 primer and Applied Biosystems Thermo Sequenase Dye Primer Manual Cycle Sequencing Kit (Thermo Fisher Scientific) according to the manufacturer’s protocol (**Fig.S2 a**).

### Gel image analysis

Each transcription product, i.e., a gel band deriving from distinct length RNA molecules (length varies because of different TSS’s) in the reaction mixture, was quantified by using Fiji software ^38^ to draw a polygonal selection contour around the gel band and calculating the total fluorescence intensity inside the contour. Background fluorescence value was subtracted from the RNA band intensity by using the average background per pixel, determined in at least 3 random RNA-free gel areas, multiplied for the contour area. The band contours in the negative control lane (no NTP’s added to the reaction) were positioned based on the RNA band positions in the NTP containing lane, and their intensities were analysed similarly. Finally, the background corrected fluorescence intensity of each RNA band was converted to molar RNA concentration by comparison to the intensity of free primer band on the lane. Since the concentration of primer was known in total sample volume at each step of in vitro transcription, the primer concentration in loaded aliquot resolved on PAGE was calculated. Because the activity of Pol I varied between different purification batches, transcription activity was normalized based on the transcription activity observed on the wild-type rDNA promoter [(−90/+30), (−45/+45) and (−30/+30)].

### Exonuclease III footprinting

DNA scaffolds for footprinting consisted of two complementary DNA strands; the strand designed to be footprinted contained Cy5.5 fluorophore at the 5’ end while the other strand was protected against exonuclease activity by connection the last six nucleotides at the 3’ end with phosphorothioates (**Table S3**) ^39^. CF·DNA complexes were formed by incubating 1.5 pmole of CF with 0.15 pmole of Cy.5.5 DNA template in DEPC-treated FTB5 buffer [50 mM Tris-HCl, 7.5, 100 mM potassium glutamate, 5% (v/v) glycerol, 5mM MgCl_2_, 0.2 mM TCEP, 0.1 mM EDTA] for 15 min at 30℃; for control reactions CF was omitted. The samples were then treated with 600 units of exonuclease III (200 U/μL stock, ThermoFisher Scientific, cat. no EN0191) for 4 min at 30℃. The reactions were stopped by the addition of 20 mM EDTA and 1% (w/v) sodium dodecyl sulphate. DNA was extracted by the additions of glycogen (RNA grade, Thermo Scientific, cat. no R0551) to final concentration of 0.5 mg/ml and chloroform:phenol:isoamyl alcohol mixture (pH 8.0, RNAse-, DNAse- and Proteinase-free, Across Organics, cat. no 327111000) followed by DNA recovery from the aqueous phase by ethanol precipitation. DNA pellet was dissolved in the 1:1 mixture of DEPC-treated water and formamide gel loading buffer, heated at 95℃ for 3 min, and then analysed on denaturing 10–12% denaturing polyacrylamide gels. Footprinting patterns were visualised with Odyssey Infrared Imager (Li-Cor Biosciences). The positions of DNA region protected by protein binding were identified by correlating the electrophoretic mobilities of the sample DNA bands with those of Sanger sequencing ladders, which were prepared similar to those for the analysis of *in vitro* transcription reactions (see above).

### Mass photometry

Mass photometry (MP) was conducted with a Refeyn Two^MP^ Mass Photometer (Refeyn Ltd) at 22°C. In the MP assay a protein molecule or complex lands on surface of the microscope slide interfering the light reflected from the surface; the change in reflected light scales linearly with the mass allowing accurate determination of molecular weights ^40^. Microscope coverslips (w×l: 24×50 mm, Refeyn) and alignment chamber (Refeyn) were sequentially cleaned with milli-Q water, isopropanol (HPLC grade) and milli-Q water followed by air stream drying. To prepare a reaction well, a silicone gasket (24 samples well cassettes for automated system, Refeyn) was then placed on the coverslip. Before each data acquisition, the z plane focus was identified using buffer-free focus method based on total internal reflection mode of the instrument. Next, 18 µL of target protein(s) or preformed CF·DNA complex was added to the reaction well and the movies containing 8853 frames were recorded for 180 s using AcquireMP software (Refeyn Ltd).

To determine the molecular masses and oligomeric states of individual proteins, stocks were diluted to 40–50 nM protein concentration in specific reaction buffers: CF (FTB5 buffer), Rrn3 (TIB2, i.e., TIB buffer with 2 mM magnesium acetate), and Pol I (TIB2). To determine the apparent dissociation constant of CF·promoter complex, 50 nM CF was mixed with 0, 25, 50, or 100 nM wild-type (WT) rDNA promoter scaffold (span -45/+45) and incubated for 5 min at 24°C before MP data collection. To study CF binding to promoters harboring a mutated base pair, 50 nM CF was mixed with 25 nM rDNA promoter (span -45/+45 and bp substitution at -27 or -28) and incubated for 5 min at 24°C before MP data collection. Finally, to evaluate holoenzyme formation, 40 nM Pol I was mixed with 80 nM Rrn3 in TIB2 buffer and incubated for 5 min at 24°C before MP data collection. Each MP analysis was performed in at least three independent experiments.

To obtain contrast-to-mass calibration of MP data, a protein standard mixture, containing BSA (molecular mass 65 kDa), IgG (150kDa monomer and 300 kDa dimer), apoferritin (443 kDa) and IgM (970 kDa) in each specific reaction buffer, was analyzed along with the actual samples in each experiment. The mean protein peak contrast was determined using Gaussian fit procedure in AcquireMP software (Refeyn Ltd). The mean contrast values from calibration protein mixture were then plotted and fitted to linear regression to define the contrast-to-mass calibration factor. MP data to determine the molecular weights and abundances of CF, Rrn3, Pol I monomers and dimers were processed, analyzed and plotted using DiscoverMP v2.3 (Refeyn Ltd). MP data to determine the apparent molecular weights and abundances of DNA-free CF and CF·DNA complex was additionally analyzed by using Origin 2016 (OriginLab Corporation) to fit the histograms of apparent molecular weight values (*m*^app^) to two-peak Gaussian equation (an area version) using *n* fixed as 2 (**Equation 1**). The fit parameters *m*_C*_, *w* and *A* are the center, width and area of the two Gaussian distributions, respectively.

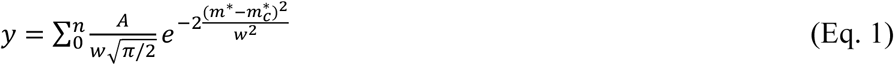

We measured blank 50 nM CF samples before each promoter binding experiment to verify the integrity CF sample. This data indicated that a minor mass peak (at about 284.4 kDa), which represented 4.3% of DNA-free CF abundance and most likely is a contaminating protein, overlapped with the mass of CF·DNA complex. This contribution was subtracted from the abundance of CF·DNA complex (*A*_CF_·_DNA_) before calculating the relative amounts of DNA-free CF and CF·DNA complex, respectively.

### Protein induced fluorescence enhancement assay: spectrofluorometer detection

The apparent equilibrium dissociation constant (*K*_Dapp_) of CF·promoter complex in manual mixing experiments was first determined for 25 nM DNA(Cy3) [contains -40/-10 rDNA promoter region and Cy3 label at the 3’ of non-template DNA (ntDNA) strand] in 50 µL of binding buffer into a quartz cuvette (Hellma Analytics, light path 3×3 mm, cat.no. HL105-251-15-40). Fluorescence intensity of this protein-free promoter sample was continuously measured for 120 s with a LS-55 fluorescence spectrofluorometer (Perkin Elmer) at 24°C. The excitation and emission wavelengths were set to 545 nm and 565 nm, respectively, and emission and excitation slits to 10 nm. The reaction mixture in the cuvette was then supplemented with 12.5–200 nM CF, thoroughly mixed, and fluorescence recording was continued for an additional 300 s. The promoter concentration after the CF addition was 24 nM. *K*_Dapp_ was estimated by fitting **Equations 2–4** to the measured fluorescence values (*F*_total_). Other parameters in **Equations 2–4** are the fluorescence intensity of protein-free DNA(Cy3) (*F*_DNA(Cy3)_), the fluorescence intensity of CF·DNA(Cy3) complex (*F*_CF·DNA(Cy3)_), the total concentration of CF ([CF_total_], the *x* axis of **Fig.2a** and **Fig.S4c**), the total concentration of labelled DNA ([DNA(Cy3)_total_]), the fraction of DNA(Cy3) bound to CF ([CF·DNA(Cy3)]_fraction_), and the fraction of protein-free labelled DNA ([DNA(Cy3)]_fraction_). Data fitting was performed using Origin 2016 (OriginLab Corporation). To determine *K*_Dapp_ from mass photometer data (label-free DNA used in MP experiments), same **Equations 2–4** were used with the exception that parameters *F*_DNA(Cy3)_ and *F*_CF·DNA(Cy3)_ in **Equation 3** were fixed as 0 and 1, respectively.

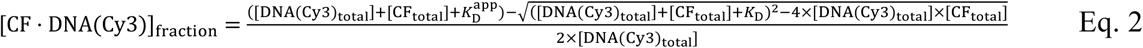

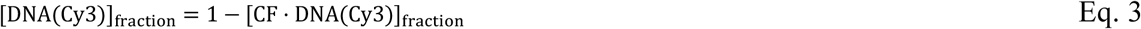

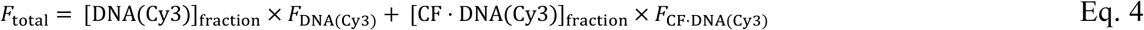

The binding of CF to different transcription DNA scaffolds was assessed in competition experiments using different label-free DNA contructs to trigger the dissociation of CF from DNA(Cy3). In competition experiments the fluorescence intensity traces were continuously measured in total for 1800 s in three sequential steps: (1) free DNA(Cy3), (2) the formation of CF·DNA(Cy3) complex, and (3) competitor-induced dissociation of CF from CF·DNA(Cy3) complex. Fluorescence intensity of 25 nM DNA(Cy3) and DNA(Cy3)·CF complex (sample contained 24 nM DNA and 120 nM CF) was determined as described in *K*_Dapp_ experiment. After the addition of competitor DNA to 227 nM concentration, the reaction mixture at the third step contained also 23 nM DNA(Cy3) and 115 nM CF. The fluorescence values were corrected for dilution accompanying the competitor addition (∼4% volume increase). The reported fluorescence dissociation traces are averages along with SE. Presented data were normalized from 0 to 1 intensity scale by setting the increment in fluorescence intensity between protein-free DNA(Cy3) and CF mixture with DNA(Cy3) (before the competitor addition) as 1 respectively. In addition to presenting data as full kinetic dissociation traces, fluorescence intensity over the last 15 s was averaged to construct bar-error graphs (average±SE) that represent the remaining fraction of CF·DNA(Cy3) complex. Data for *K*_Dapp_ estimation and competition of label-free DNA templates was measured in FTB5 buffer at least in three independent experiments and averaged. *K*_Dapp_ at reduced buffer ionic strength was determined in LTB5 [25 mM Tris-HCl, 50 mM K-glutamate, 5% glycerol, 5 mM MgCl2, 0.1 mM TCEP, 0.05 mM EDTA].

### Protein induced fluorescence enhancement assay: stopped-flow detection

Measurements were performed at 24°C using SFM-3000 stopped-flow (SF) instrument equipped with FC-08 reaction cuvette (BioLogic). The instrument was operated using 4 ms sample mixing dead time. Cy3 fluorophore in the promoter templates was excited at 553 nm and emitted light was collected through a 570 nm longpass filter. Total reaction volume in SF experiments was 150 µL, which was obtained by mixing 75 µl from each of two active sample storage syringes (one typically containing CF and the other containing the promoter sample) in the SF instrument.

The rate of CF binding to the promoter DNA was determined by mixing to final concentrations 12.5−120 nM CF with 24 nM DNA(Cy3) scaffold in FTB5 buffer. Fluorescence signal (i.e., detector voltage) of the reaction mixture was continuously recorded for 100 s using two time-base settings: for reaction time range 0.004–7.45 s data was integrated 0.5 ms per each time-point while for the range 7.47–100 s the integration was 50 ms. Fluorescence signal of protein-free DNA was measured by mixing DNA(Cy3) with FTB5 buffer.

The rate of CF dissociation from the promoter DNA was determined by mixing preformed CF·DNA(Cy3) complex [concentrations after mixing: 115 nM CF and 23 nM DNA(Cy3); twofold higher during the preformation] with 227 nM (after mixing) unlabelled competitor DNA promoter (−30/30). Reference signal of protein-free DNA(Cy3) promoter was determined by mixing 23 nM DNA(Cy3) and FTB5 to the reaction cuvette of the SF instrument. Fluorescence intensity was recorded using 0.5 ms and 50 ms data integration times for reaction time ranges 0.004–7.45 s and 7.47–250 s, respectively.

Each reported SF curve is the average of 6–8 individual SF traces (a.k.a mixing reactions or SF shots) from which the fluorescence curve of protein-free DNA(Cy3) has been subtracted, and which was followed by the normalization of the initial fluorescence level of the reaction (over the first ∼0.3 s where CF binding/dissociation is not yet observed) as 1. For CF dissociation data the reaction end-point was additionally normalized to zero fluorescence. The apparent rate of CF binding to promoter DNA (*k*_obs_) was extracted by fitting **Equation 5** to fluorescence data: *A* in the equation is the fluorescence change amplitude of the exponential reaction phase, *y*_0_ is the fluorescence at the end-point of the exponential reaction phase, *t* is the reaction time (i.e., abscissa), and *m* is the slope of the linear reaction phase. CF binding data obtained using wild-type DNA(Cy3) was strictly mono-exponential whereas mutant scaffolds [DNA(Cy3,-27) and DNA(Cy3,-28)] had an additional slow linear increase in fluorescence. Consequently, *m* was set as 0 for wild-type data and left as a free parameter for mutant data.

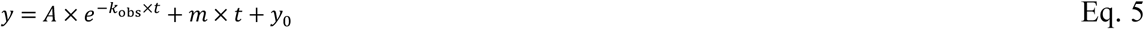

The median rate of CF dissociation from promoter DNA (*k*_rev_) was extracted by first fitting the stretched mono-exponential function (**Equation 6**) to fluorescence curves ^41,42^ and then calculating *k*_rev_ using **Equations 7** and **8**. The stretching parameter *β* modifies the product of the rate parameter (*k*) and reaction time (*t*) to accommodate the deviation of the reaction progress curve from single exponential behaviour (see also Results section). Data fitting was performed using Origin 2016 (OriginLab Corporation).

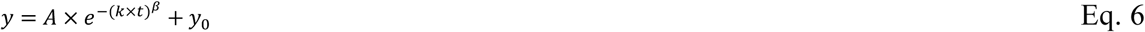

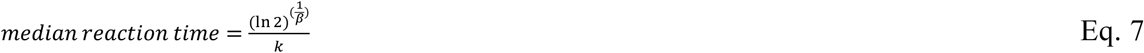

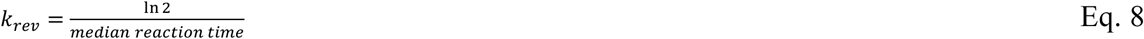

### Molecular modelling

System preparation. All modelling was conducted with Schrödinger Suite software version 2024-1 (Schrödinger Inc.). The starting structure for the modelling was cryo-EM based structure of *S. cerevisiae* preinitiation complex, showing Pol I, Rrn3 and CF bound on a rDNA promoter at 2.90 Å resolution ^11^. This structure (‘RNA Polymerase I closed conformation 2’; accession code: 6RQL) was downloaded from the PDB database and prepared using the Protein Preparation Workflow of Maestro with default settings. This included filling in missing sidechains, optimizing hydrogen bonds and protonation using PROPKA at pH 7.4, deleting water molecules >5Å from the heteroatoms and a short energy minimization using OPLS4 force field ^43^. The initial structures for the complexes with DNA mutations were constructed from this pre-processed structure by manually mutating the base pairs at position -28 or -27 of the rDNA promoter from C·G to A·T (tDNA·ntDNA) and from A·T to C·G, respectively.

Simulations. For the molecular dynamics (MD) simulations, the complex was solvated with SCP water molecules with 10 Å buffer distance and the system was neutralized by adding 61 sodium ions. The resulting simulation systems had ∼740 000 atoms. MD simulations were conducted with Desmond using default parameters from Schrödinger Suite version 2024-1 on the CSC (IT Center for Science, Finland) supercomputer Puhti. The default relaxation protocol of Desmond was used before the 80 ns production simulations, which were conducted in NPT ensemble (temperature: 300 K; thermostat: Nosé–Hoover chain; pressure: 1.01325 bar; barostat: Martyna–Tobias–Klein) using the OPLS4 force field. The default time step of 2 fs and cutoff radius of 9.0 Å for coulombic interactions were used. The total simulation time was 240 ns (3 × 80 ns) for all systems and the frames were saved for every 0.2 ns. For analysis, the simulations were concatenated using trj_merge.py script and aligned using trj_align.py script built-in in Schrödinger Suite. The aligned trajectories are available at Zenodo in .cms format (DOI 10.5281/zenodo.14001801).

## RESULTS

### Oligomeric state of purified RNA polymerase I and transcription initiation factors

To investigate the mechanism by which transcription initiation factors and Pol I identify the rDNA promoter and TSS, we reconstituted yeast Pol I *in vitro* transcription system for functional studies using purified proteins and nucleic acids. To this end, we expressed *Saccharomyces cerevisiae* transcription initiation factors CF and Rrn3 in *Escherichia coli* and purified the recombinant proteins using several chromatographic techniques (**Fig. S1a,b**). Pol I was purified from *S. cerevisiae* cells harvested in the exponential growth phase (**Fig. S1c**).

The oligomeric state of transcription factors (TFs) was initially assessed by inspecting the chromatograms of the gel-filtration (GF) purification step. CF emerged from the column in a single major peak, suggesting only one oligomeric state for this protein. (**Fig. S1d**). In contrast, Rrn3 was present in two well-separated chromatogram peaks, presumably representing its two distinct oligomers (**Fig. S1e**). We then utilized mass photometry (MP) ^40^ to directly determine the molecular weights of single protein molecules/complexes present in the Rrn3 and CF GF eluates in buffer compositions similar to those used in transcription *in vitro*. The mass distribution histogram of Rrn3 GF monomer preparation (p4 in **Fig.S1e**) elucidated the co-presence of Rrn3 monomers (84% of all molecules) and dimers (16%) in the sample indicating dynamic equilibrium between monomer and dimer oligomeric states (**Fig. S1f**). The experimentally determined molecular weights of the monomer (82.3±12.9 kDa) and dimer (145±25 kDa) Rrn3 are consistent with the masses calculated from the protein sequence (72.4 and 144.8 kDa). The molecular weight histogram of CF indicated only one significant protein population with the estimated molecular mass of 219±33.8 kDa, a value in excellent agreement with the calculated mass of CF monomer (221.8 kDa) (**Fig. S1g**). In the case of Pol I, the MP analysis indicated the main population of monomeric protein (86% of all molecules) and a minor dimeric population (13%) (**Fig. S1h**). The experimentally determined molecular weights of Pol I monomer (577±88 kDa) and dimer (1141±109 kDa) matched very well with the calculated values (589.5 and 1179 kDa). Overall, the molecular weight analysis indicated that the purified proteins were obtained in their correctly assembled states.

### Binding of CF to rDNA promoter: location, stoichiometry and affinity

CF is regarded as the key TF that recognizes the rDNA promoter and recruits Pol I to it^11,18,29^. We therefore decided to investigate, at a previously lacking quantitative level, how CF interacts with the rDNA promoter. To proceed towards this aim, it was essential to first define the boundary of CF binding site on the promoter and specificity of its selection. To achieve single base pair (bp) resolution, we employed Exonuclease III (ExoIII) footprinting assay based on the detection of Cy5.5 label at 5’ end of DNA template where the complementary strand has phosphothioate protection at 3’ end^39^. To setup the ExoIII assay, we built two synthetic DNA scaffold models of the rDNA promoter, covering the promoter region with reference to TSS at +1 from nucleotide position -45 to +45 [Cy5.5(−45/+45) and (−45/+45)Cy5.5] and from -80 to -35 [Cy5.5(−80/-35)] as a control of non-specific interaction(**Fig. 1a and Fig. S2a**).

**Figure 1.**
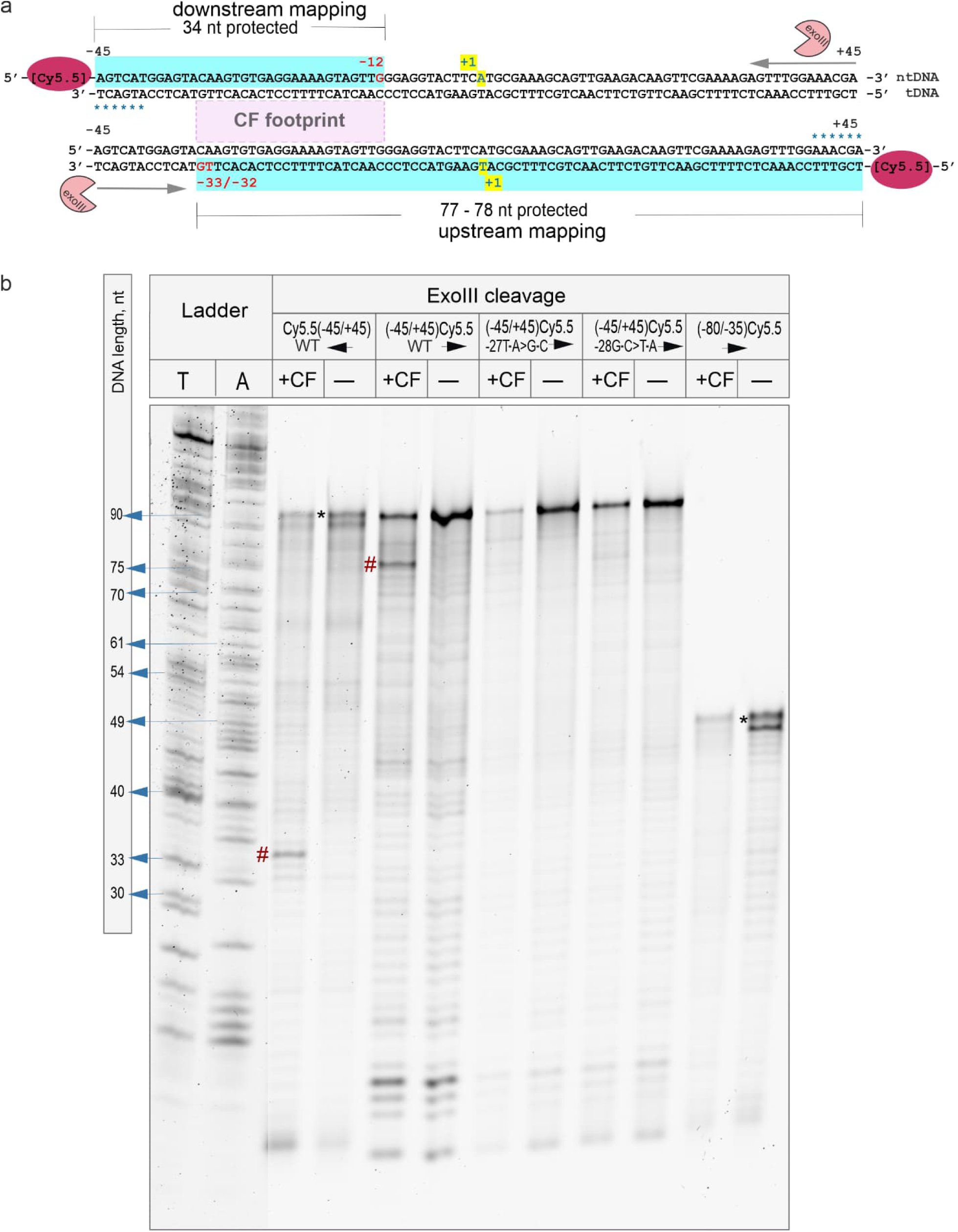
Footprinting of CF on wild-type and base substituted rDNA promoters. (a) The cartoon represents ExoIII-based footprinting of CF binding site on the wild-type rDNA promoter templates (span -45/+45, TSS at +1 highlighted with yellow). To identify the downstream and upstream boundaries of the CF footprint, the 5’end of either non-template (promoter shown on the top) or template strand (bottom promoter) is labeled with cyanine 5.5 fluorophore (red ellipses); the complementary strand was protected at the 3’ end with phosphorothioate bonds (blue asterisks). The direction of ExoIII cleavage is shown with grey arrows. The protected area on the non-template DNA (Cy5.5-ntDNA) and template DNA (Cy5.5-tDNA) strands is highlighted with cyan rectangles and the detected CF footprint is marked with the light purple rectangle. (b). Mapping CF boundaries on Pol I promoter templates by ExoIII. Gel lanes from left to right: dTTP (T) and dATP (A) ladder (Sanger sequencing), footprinting of the wild-type rDNA promoter for the downstream boundary (indicated by left black arrow) in the presence (WT, +CF) and absence (WT, -) of CF, footprinting of the wild-type rDNA promoter for the upstream boundary (indicated by right black arrow) in the presence (WT, +CF) and absence (WT, -) of CF, the upstream footprinting of rDNA promoter (− 45/+45) substituted at position -27 with (−27T·A>G·C; +CF) and without (−27T·A>G·C; -) the addition of CF addition, and at -28 position with (−28G·C>T·A; +CF) and without (−28G·C>T·A; -) the addition of CF; and the rDNA promoter fragment with span -80/-35 in the presence (NC, +CF) and absence (NC, -) of CF. Blue arrows indicate the electrophoretic mobility of specific length DNA ladder strands. Full-length promoter is indicated with asterisks and CF footprint with hash mark.

The footprinting results show that CF binds to the -45/+45 promoter but not the -80/-35 promoter. This selective binding was evident by the presence of CF-dependent protected region on the -45/+45 promoter, whereas no such protected region was observed on the -80/-35 promoter (**Fig. 1b**). PAGE analysis of the untreated scaffolds indicated excellent DNA purity, facilitating the mapping of exact footprint boundaries (**Fig. S3a**). To map the footprint, we compared the lengths of the protected DNA strands to a DNA ladder of defined lengths and found the CF to protect the rDNA promoter fragment from -32 to -12 (**Fig. 1**) (**Table S4**). Our interpretation aligns with the previously determined CF footprint (from the position -32 to -9 bp)^11^, the span of the minimal binding competent DNA fragment (−28 to -17 bp)^34^ and structural evidence from cryo-EM models (−35 to -12 bp^9^, -40 to -16 bp ^10^). We were further interested in the stability and possible DNA unwrapping–wrapping dynamics in the CF·promoter complex, and therefore studied the time-course of ExoIII digestion (**Fig. S2b**). The downstream edge of the footprint remained sharp at position -12 in all time-points indicating that DNA at the downstream part of the CF binding site remains tightly anchored to CF with non-detectable level (if any) of unwrapping dynamics. The close inspection of the upstream CF footprint reveals two bands, suggesting that most CF·DNA complexes have the footprint edge at -32 while a smaller fraction of the complexes have the edge at -33. Overall, CF protects invariable DNA region over the increasing cleavage time, indicating strong and specific CF binding to the promoter DNA (**Fig. S2b**).

To track directly the binding dynamics of CF, we constructed a DNA scaffold containing the promoter sequence from -40 to -10, which encodes the CF binding motif determined by footprinting analysis and has a cyanine 3 (Cy3) fluorophore attached at the downstream end of the DNA fragment [hereafter called as DNA(Cy3)] (**Fig. S4a**). Cy3 was selected due to its tendency to exhibit a significant increase in fluorescence intensity upon protein binding to a nearby site on the DNA ^44^. This amplification in Cy3 intensity stems from its cis/trans photoisomerization, commonly known as protein-induced fluorescence enhancement (PIFE). Indeed, upon the addition of CF to DNA(Cy3) sample, the fluorescence intensity increased by ∼35% (**Fig. S4b**). This PIFE effect was reversed when the dissociation of CF from DNA(Cy3) was triggered by the addition of unlabeled rDNA promoter to the sample. These observations suggest that CF binds near Cy3, consistent with the orientation of CF binding sequence motif on the DNA scaffold. We first estimated the binding affinity of CF to DNA(Cy3) using a spectrofluorometer to record changes in Cy3 fluorescence and manual reagent addition steps. The obtained equilibrium binding curve was described by a simple model, which assumes that one CF molecule binds to one DNA(Cy3) molecule (**Fig. 2a**). The corresponding fit equations (**Eq. 2–4**) additionally take into account changes in free CF concentration upon DNA binding. The binding affinity was determined in two buffers conditions; FTB5 buffer has a moderate ionic strength of 156 mM while LTB5 has it lowered to 85 mM. The apparent dissociation constant (*K*_Dapp_) was found to be 106±64 nM (standard error, SE) of CF in FTB5 (**Fig. 2a**) and twofold smaller, 52±31 nM, in LTB5 (**Fig. S4c**). All fit parameters are shown in **Table S5**. Based on the binding inhibition by increased ionic strength, it appears that ions and water molecules contribute to the association of CF with the specific promoter binding site, as previously analyzed in-depth for the interaction of *lac* repressor with DNA ^45^.

**Figure 2.**
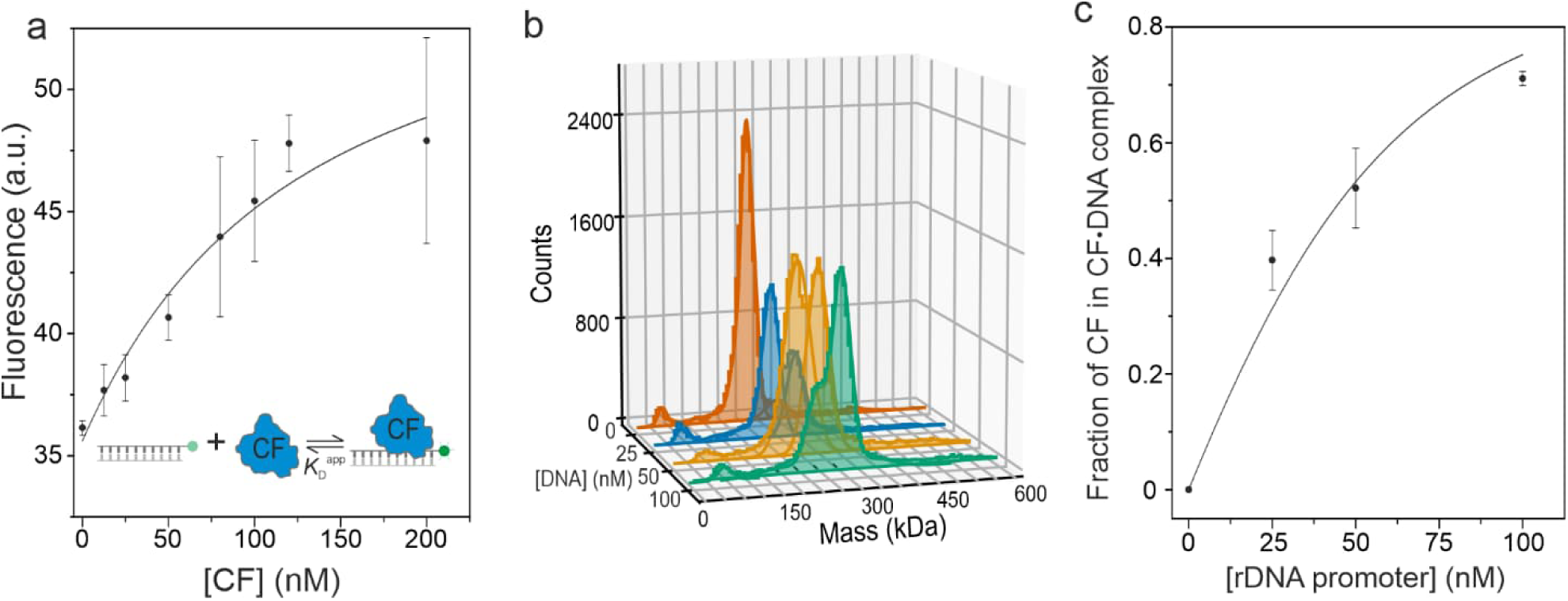
Binding affinity of CF to rDNA promoter. (a) Apparent dissociation constant of CF·DNA(Cy3) complex (*K*_Dapp_) was determined by mixing 24 nM DNA(Cy3) promoter with 12.5– 200 nM CF. Fluorescence intensities were measured using spectrofluorometer. Data points and error bars are averages and standard deviations from 3–7 independent experiments, respectively. (b) Mass photometer based mass distributions of free and DNA-bound CF molecules/complexes. Samples contained 50 nM CF and variable concentration (0, 25, 50, or 100 nM) of unlabeled rDNA promoter (span -45/+45). The solid lines over the distributions indicate the best Gaussian fits of molecular masses (kDa). (c) The relative amount of CF bound to rDNA promoter in MP assay (n=3 independent experiments). Theoretical curves in panels a and c were obtained by fitting Equations 2–4 to data.

To further dissect the stoichiometry of CF·promoter complexes, we employed mass photometer to compare the molecular mass distribution of CF (50 nM) in the presence of 0–100 nM rDNA promoter (span -45/+45). MP data revealed promoter-dependent formation of a new protein population at 275±12.2 kDa (n=9) and a corresponding decrease in the abundance of DNA-free CF monomer at 228±6.3 kDa (n=9) (**Fig. 2b**). We interpret the appearance of population at 275 kDa following the addition of promoter DNA as the CF·promoter complex with 1:1 stoichiometry (the calculated mass of the promoter is 55 kDa), consistent with PIFE data. To estimate *K*_Dapp_ from MP data, we first corrected the CF-promoter abundance for the apparent protein impurity, which was present as a minor factor (6.4%) in DNA-free CF sample and whose mass (284±11.3 kDa) overlapped with CF·DNA complex. Fitting of **Eq. 2–4** to data estimated *K*_Dapp_ as 20.5±4.5 nM (SE) (**Fig. 2c**, **Table S5**). Noteworthy, the binding affinity (*K*_Dapp_) reported by PIFE and MP experiments are not necessarily identical. The PIFE assay was designed to report the binding of CF to its specific binding site near Cy3 label, while MP can additionally detect unspecific CF·DNA complexes or the intermediate complexes on the specific binding pathway, provided that those complexes are sufficiently stable not to dissociate within the 0.02 s data frame in the recorded MP movies.

### Binding of CF to rDNA promoter: kinetics

To determine the rate of CF binding to the rDNA promoter, we employed a stopped-flow (SF) instrument to automate the mixing of CF from one syringe with DNA(Cy3) (−40/-10 fragment) from the other syringe, thereby reducing dead time to 4 ms. We performed the SF experiment at several CF concentrations (12.5–120 nM), keeping the DNA(Cy3) concentration fixed at 24 nM. The reaction progress curves started to show a noticeable increase in Cy3 fluorescence intensity in about 1 s after mixing the DNA(Cy3) with CF, the intensity reaching maximum by about 80 s (**Fig. 3a**). We calculated the change in the Cy3 intensity in each CF concentration and fit **Eq. 2–4** to this data to obtain *K*_Dapp_ as 79 ± 35 nM (SE) (**Fig. 3b**), a value consistent with *K*_Dapp_ from manual PIFE experiments (**Fig. 2b**).

**Figure 3.**
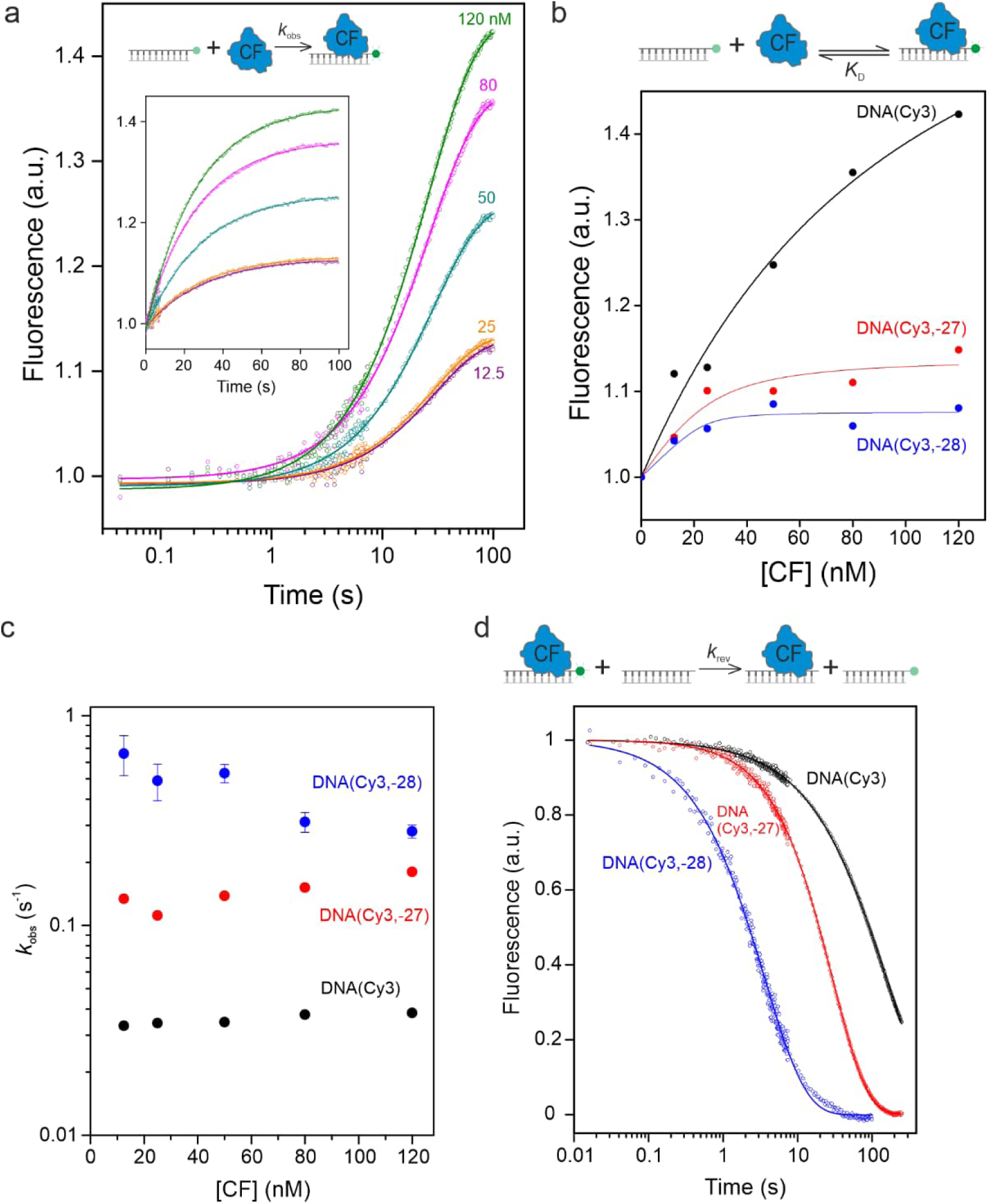
Interaction dynamics of CF and rDNA promoter. (a) SF fluorescence traces monitoring the binding of 12.5–120 nM CF to 24 nM wild-type DNA(Cy3) are shown. The fit curves were obtained using Eq. 5. The inset shows same data with linear timebase. (b) The fluorescence intensity of WT or mutant DNA(Cy3) scaffolds at the end of CF binding reaction is shown based on the recorded intensity 100 s after the mixing of CF and DNA. The fit curves were obtained using Eq. 2–4. (c) The observed rate constants (*k*_obs_) of CF binding to WT or mutant DNA(Cy3) scaffolds. (c) The rate of CF dissociation from WT or mutant DNA(Cy3) scaffolds was examined by challenging preformed CF·DNA(Cy3) complex with excess of label-free promoter DNA (span -30/+30). The fit curves were obtained using Eq. 6.

To initially visually compare the binding rates in different CF concentrations, we equalized all signal amplitudes (**Fig. S5a**). The normalized curves reveal that the reaction half-life (i.e. the time-point in which half of the fluorescence increase happened) was similar, approximately 16–20 s, in all CF concentrations. It thus appears that the apparent reaction rate is not determined by a bimolecular binding reaction, i.e. the combination of free CF and free DNA(Cy3) to a binary complex, and is instead determined by a unimolecular reaction taking place in the preformed CF·DNA complex, e.g., a conformational reorientation of CF on the DNA. To obtain quantitative rate parameters, we fit the original stopped-flow reaction progress curves to a single exponential rate equation (**Eq. 4**) (**Fig. 3a**). The obtained fit parameters confirm that that the observed rate of CF·DNA complex formation (*k*_obs_) has no significant dependence on the CF concentration; specifically, the *k*_obs_ values were found as 0.033–0.038 s^-1^, the average *k*_obs,avg_ as 0.036±0.002 s^-1^ and the average reaction half-life as 19.5±1.1 s [calculated by ln(2)/*k*_obs,avg_] (**Fig. 3a**, **Table S5**).

The binding of CF to its cognate binding site on the DNA is a reversible reaction. Such a reversible system approaches equilibrium at an observed rate, which is defined by the sum of forward and reverse reaction rates (*k*_obs_=*k*_for_+*k*_rev_;)^46^. To determine the dissociation rate, we conducted sequential addition experiments, in which CF was first mixed with DNA(Cy3) to form CF·DNA(Cy3) complexes and in which the addition of excess unlabelled competitor rDNA promoter in the second step triggered the breakage of CF·DNA(Cy3) complexes (**Fig. 3d**). The preliminary analysis of CF dissociation time traces revealed that a fit using the single exponential rate equation yielded poor results, as evident from the systematic deviations of the trend curve (**Fig. S5b**). To obtain a good fit, we employed a stretched exponential function (**Eq. 6**), an empirical function often used to describe heterogeneous systems, including enzyme reactions ^41,42^. Based on a previous discussion of different potential sources of heterogeneity,^42^ we assume that in our case the stretched exponential fit, specifically the stretching parameter *β*, accommodates both temporal and structural heterogeneity of CF·DNA complexes, as well as deviations from single exponential behavior caused by the sequential nature of the dissociation reaction (note that also the binding happened in a two-step reaction). For a similar system a meaningful interpretation of the rate parameter (*k*) was previously obtained by first calculating the median reaction time (**Eq. 7**) and then converting it to the median reaction rate (**Eq. 8**).^42^ The fit parameters shown that CF dissociates slowly from its specific binding site on the DNA with the median reaction time 93.6±2.1 s and the median dissociation rate (*k*_rev_) 0.0076±0.0002 s^-1^ (**Table S5**). For comparison, data analysis using the single and double exponential rate equations returned dissociation rates of 0.0112±0.0001 s^-1^ and 0.0084±0.0001 s^-1^ (the fit assigned 88.5% of total reaction amplitude to this rate), respectively, indicating that the inferred dissociation rate is only slightly affected by the selected fit equation (**Fig. S5b**). Finally, *k*_for_ was calculated (from *k*_obs_=*k*_for_+*k*_rev_), using known *k*_obs_ (0.036±0.002 s^-1^) and *k*_rev_ (0.0076±0.0002 s^-1^), to be 0.0284±0.0022 s^-1^. Overall, these parameters indicate that both the formation and dissociation of the specific CF·DNA complex is limited by slow isomerization steps, the reaction towards dissociation direction (*k*_rev_) being 3-fold slower than to the forward direction (*k*_for_).

### Recruitment of Pol I to rDNA promoter

CF recruits Pol I to the rDNA promoter during the formation of preinitiation complex (PIC). We sought to evaluate the efficiency and stability of this recruitment by ExoIII footprinting the rDNA promoter from the downstream direction after initiating PIC assembly on promoter. The results indicate that the edge of the protein-protected promoter DNA fragment locates at position -12 and is thus indistinguishable from the downstream footprint created by CF binding alone (**Fig. S3b**). The same CF footprint was also obtained in the presence of NTPs (A+U±G) supporting the synthesis of 2 and 3 nt long RNAs in abortive initiation reaction (**Fig. S3c**). The lack of Pol I footprint in the absence of NTPs indicates that CF cannot recruit Po1 stably to the rDNA promoter. The lack of Pol I footprint in the abortive initiation conditions further indicate that the open complex, i.e., the conformation of PIC containing a melted transcription bubble, is also unstable. Because we observed robust RNA synthesis at the promoter by Pol I in the presence of NTPs and CF but not in the absence of CF (**Fig. 4a**, see also the inset), CF can recruit Pol I to the rDNA promoter, but this recruitment is likely short-lived before Pol I dissociates.

**Figure 4.**
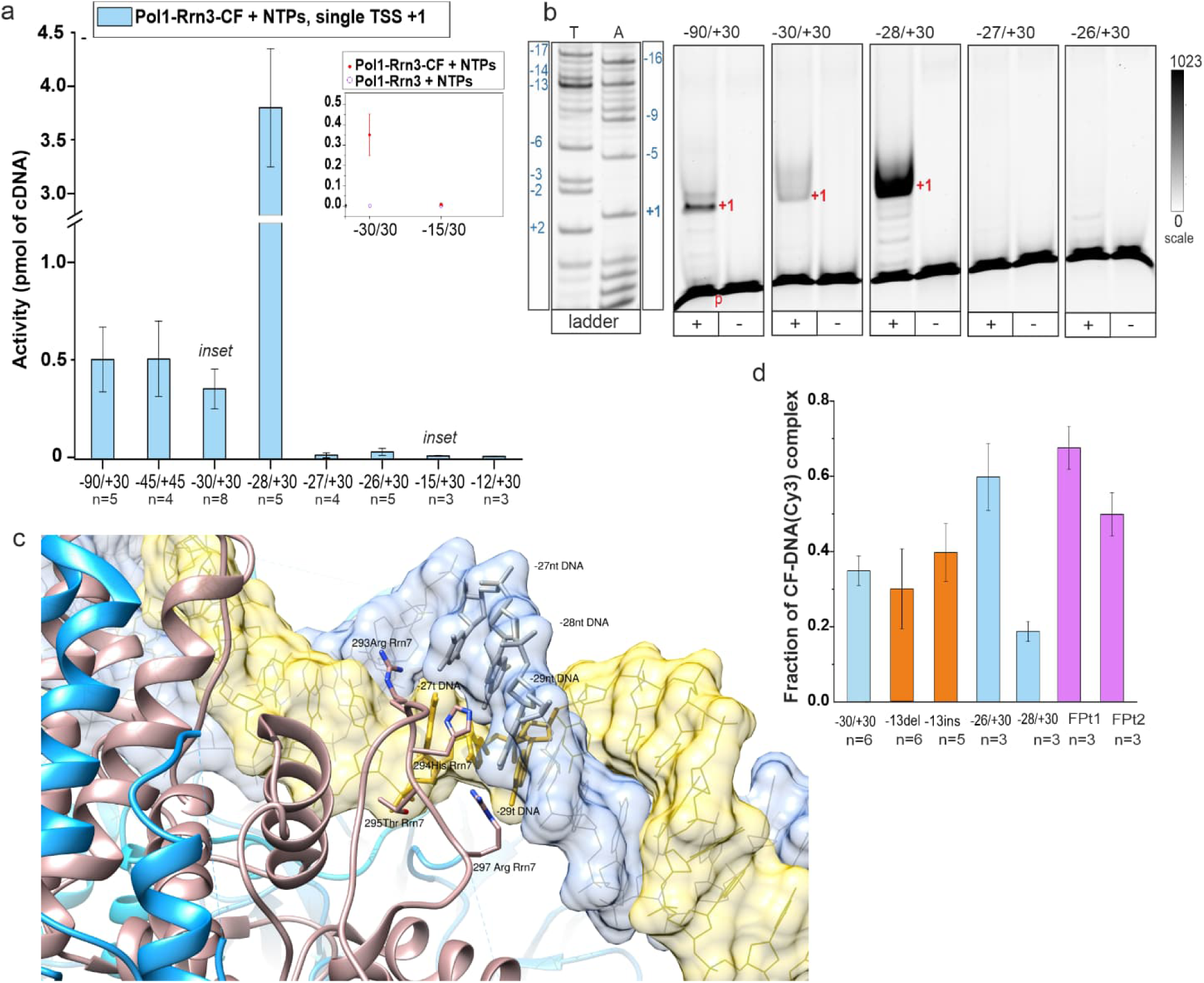
Role of CF and promoter CF-binding site in transcription initiation. (a) Transcription initiation activity on different length (span -xx/+yy) WT promoter are shown. The transcription reaction contained rDNA promoter, Pol I, Rrn3, CF, and NTPs. All active promoters had the transcription start site (TSS) at position +1, the cDNA products of rRNA transcripts were quantitated and plotted. Data is average activity and SD (error bars) from the indicated number (n) of independent experiments. The inset demonstrates the transcription activity in the presence and absence of CF. (b) Example of PAGE gel illustrates the TSS position and the level of transcription activity as detected by primer extension assay. T and A indicate Sanger sequencing ladders prepared using TTP and ATP analogues as chain terminators, respectively; the promoter positions corresponding to different ladder bands are indicated in blue numbers. All reactions presented in the figure were performed simultaneously and resolved on the same gel. Red numbers and p indicate the inferred TSS’s and the primer, respectively. All assembled PIC’s were split to transcription reaction supplemented with NTPs (indicated with +) and negative control without NTPs (−). (c) Rrn7 subunit of CF (colored in brown) embraces rDNA promoter near the positions -27 and -28. Several residues (Arg293, His294, Arg297) orient into the promoter major groove of promoter core element potentially enabling affinity of CF via electrostatic attraction to the negatively charged DNA backbone or base-specific interactions. Polar Thr295 is also found in the major groove. Polar residues are frequently acting as both hydrogen bond donors and acceptors in their interaction with nucleobases with a preference for the adenine ^69^. Rrn6 subunit of CF is represented in meadow blue. Nucleotides at positions -27, -28 and -29 are highlighted: in yellow in the tDNA and in silver in the ntDNA. DNA surface is colored in semi-transparent shades: pale yellow for ntDNA and sky blue for tDNA. The figure was produced using cryo-EM model 6RQL and Chimera 1.14 software ^70^. (d) Fractions of CF·DNA(Cy3) complex were determined following the incubation with label-free DNA competitors in the SF instrument. Linear (−30/+30; -28/+30; -26/+30), single base pair mutated (−13 del; -13ins) and fork (FPt1; FP2) promoters were used as competitors. The fluorescence intensity of protein-free DNA(Cy3) and CF·DNA(Cy3) complex were set as 0 and 1, in each measurement respectively. Data is average and SE from the indicated number (n) of independent experiments.

### Promoter sequence determinants of transcription initiation efficiency

We next assessed the promoter sequence determinants of the transcription initiation efficiency, focusing specifically on the region within and around the CF binding site. We first aimed to define the upstream edge of the functional promoter and evaluate whether promoter elements beyond it contribute to the transcription initiation efficiency. We therefore examined the transcription efficiency of the basal system, consisting of Pol I, Rrn3 and CF proteins, on a series of rDNA promoter scaffolds, which all had the downstream end fixed at +30 but differed with respect to their upstream edge/length. Our data indicates steady transcription rate when the upstream end of the promoter is trimmed from -90 to the border of CF binding site at -30 (**Fig. 4a**,**b**). The finding suggests that the basal transcription system cannot utilize the promoter region upstream from the CF binding site to tune the initiation efficiency, contrasting Pol I from bacterial RNA polymerase, which interacts with and bends the promoter upstream to facilitate the formation of the open complex and rate of transcription initiation.^47,48^ As expected, the deletion of CF binding site, by placing the promoter upstream end to -15 or -12, abolished all activity. The partial deletion of CF binding site, by placing the promoter upstream end to -26 or -27, also inactivated transcription. We mapped the critical upstream border of the promoter to -28 as this promoter supported strong activity that in fact was 7.5-fold more than when the promoter contained more extended upstream sequence. Noteworthy, the cryo-EM based model of Pol I preinitiation complex (PDB access code: 3RQL)^11^ indicates the widening of the DNA major groove from the positions -22 to -27 in comparison to standard B-DNA, probably because of the insertion of the C-terminal cyclin fold of Rrn7 (subunit of CF) and the N-terminal DNA-binding helical bundle of Rrn11 (subunit of CF) into this groove (**Fig. 4c**). It is thus plausible that the removal DNA upstream from -27 reduces the energetic penalty of the DNA major groove widening, leading to stronger CF binding and higher transcription initiation activity.

Further visual inspection of the cryo-EM model reveals that Rrn7 residues H294 and R293 locate adjacent to base pairs -27/-28 and -26/-27, respectively, implying possible sequence specific recognition of the base or base pair identities in these positions (**Fig. 4c**). To understand the role of single base pairs at the upstream edge of the CF binding site, we substituted either base pair -27 (T·A>G·C, non-template DNA·template DNA) or -28 (G·C>T·A) and determined the effect on the transcription initiation activity (**Fig. 5a**). The substitution at -27 caused about tenfold decrease in the activity while the substitution at -28 decreased the activity below detection limit (**Fig. 5b**). The specific interaction of CF with the base pairs at positions -27 and -28 appears thus essential for the positioning of CF on the rDNA promoter in a manner that supports the recruitment of Pol I.

**Figure 5.**
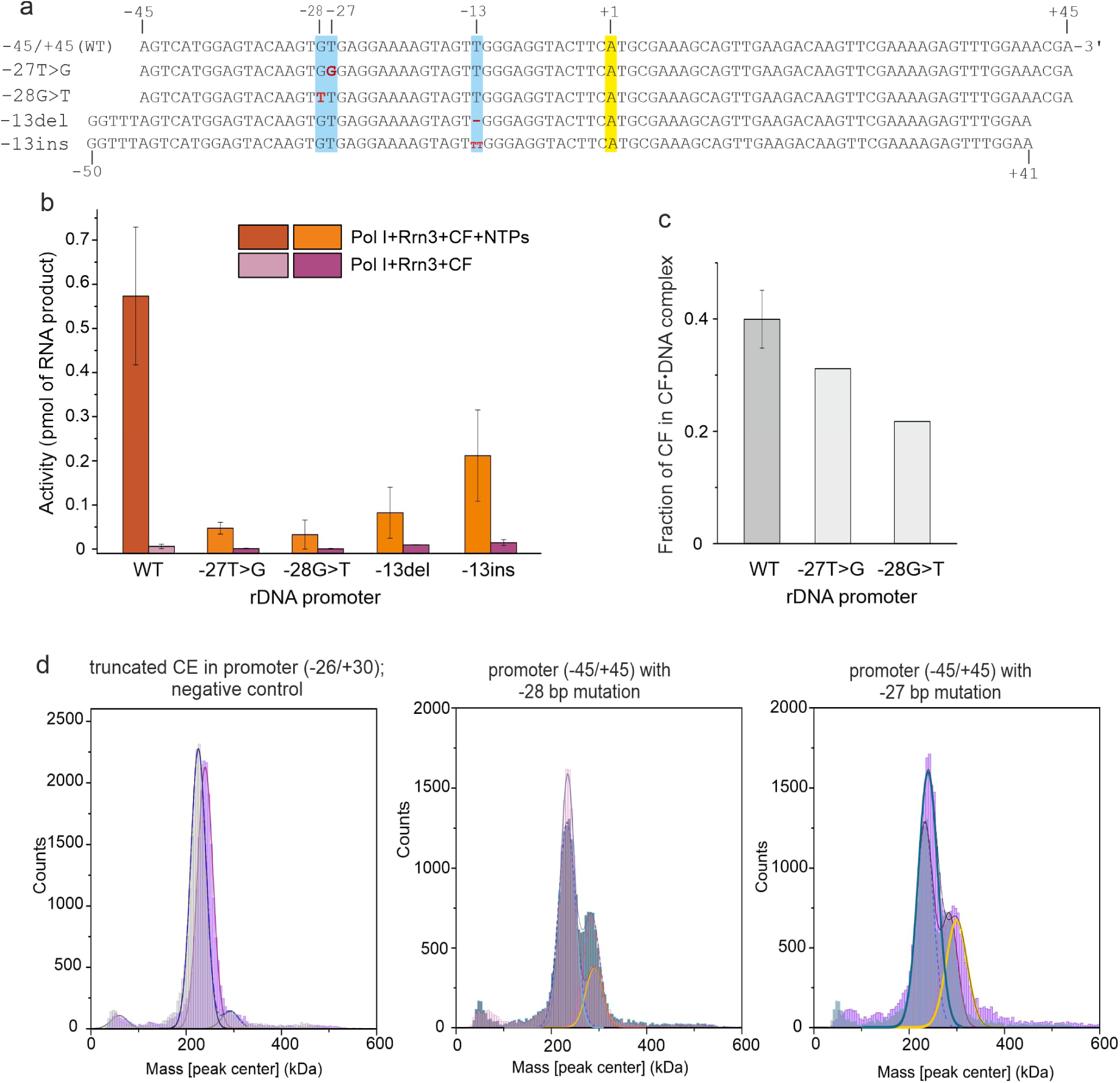
The effects of single base pair mutations on the promoter recognition and transcription activity. (a) Non-template DNA strand sequences of rDNA promoter scaffolds incorporating base pair mutations used in the transcription activity and mass photometry based CF binding assays are shown. The locations of base substitution and insertion/deletion are highlighted with light blue shading and specified in red font. The TSS is highlighted with yellow. The numeric position of these sites are indicated on top of the sequences. (b) Transcription initiation activity of Pol I on WT and mutant rDNA promoters is shown as averages and SE of three independent experiments. (c) The fraction of CF bound to rDNA promoters was determined using MP. 50 nM CF and 25 nM promoter concentrations were used. (d) Histograms of CF mass distributions after the addition of (−26/+30) truncated promoter (left panel) and promoter mutated at -28 bp (middle panel) and -27 bp (right panel) respectively are colored in lilac. The mass distribution of CF sample with the truncated promoter (−26/+30) is overlayed with the mass distribution of free CF, which is shaded in grey in the left panel. The mass distributions of CF samples with point-mutated promoters are overlayed with the mass distribution of CF sample with ET promoter (−45/+45), which is shaded in blue in the middle and right panels.

The comparison of different cryo-EM reconstructions suggest that CF and the upstream DNA can pivot around Pol I up to 15°, thereby potentially driving DNA melting inside Pol I.^10,11,49^ To gain functional insight on how delicate the positioning of CF and Pol I and its dynamics on the promoter are, we either deleted the base pair at position -13 (−13del promoter) or inserted an additional base pair between the original positions -13 and -12 (−13ins promoter) in the linker region of the rDNA promoter (**Fig. 5a**). If the protein system would respond rigidly without any rearrangement, these promoter DNA mutations would simultaneously change the contact distance and angle of CF and Pol I by ±3.4 Å (±1 bp) and ±36°. We observed that both mutations caused the suppression of transcription initiation activity; the deletion and insertion of a bp decreasing the transcription level by 85 % and 65%, respectively (**Fig. 5b**). Neither of the mutations affected the transcription start site, the RNA synthesis initiating consistently from +1 (**Fig. 5b**). These finding indicate that the proper spacing of CF and Pol I binding sites on the promoter is important but the proteins can adjust and partially compensate for small changes (here 1 bp indel), probably because the interaction surface of CF with Pol I is relatively small and, in large part, formed by protein domains, specifically Zn-ribbon and B-reader in Rrn7, at the end of flexible B-linker structure^11^. The invariability of TSS further imply that Pol I, at least to a limited degree, senses the promoter sequence or structure to identify the correct TSS instead of solely relying on the CF-mediated recruitment on the DNA.

### Effects of binding site mutations on the interaction with CF

To gain quantitative insight into the effects of DNA major groove mutations on CF binding, we reproduced the substitutions at positions -27 or -28 in DNA(Cy3) scaffold (span: -40/-10, Cy3 fluorophore at the 3’ end of ntDNA) to create mutant rDNA promoter scaffolds DNA(Cy3,-27) and DNA(Cy3,-28) for the PIFE-based binding studies (**Fig. S4a**). We then mixed 24 nM DNA(Cy3,-27) or DNA(Cy3,-28) with different concentrations of CF in the SF instrument and calculated the change in the Cy3 intensity (**Fig. 3b**). CF binding caused significantly smaller intensity increase on both mutant scaffolds in comparison to wild-type DNA(Cy3). This effect, in principle, is expected if the DNA mutations drastically decrease the overall binding affinity of CF. However, the CF concentration dependence of PIFE saturated within the used CF concentrations (up to 120 nM) as indicated by the estimated *K*_Dapp_ values 7±7 nM [DNA(Cy3,-27)] and 1±3 nM [DNA(Cy3,-28)] (**Table S5**). It thus appears that the DNA major groove mutations compromised the capability of CF to recognize its specific binding site (near Cy3) on the mutant scaffolds, leading CF to be mostly bound at the unspecific binding sites, which are distributed along the DNA scaffold, do not produce PIFE effect, and cannot be overcome by increasing CF concentration in the experiment. To corroborate this model, we analyzed CF binding to -27 or -28 mutated promoters using mass photometer, which may detect both specific and unspecific CF·DNA complexes. Data indeed indicates significant CF binding occupancy on the mutant promoters, specifically about 75% (−27 mutation) and 50% (−28 mutation) of the WT binding level (**Fig. 5c**). Also consistent with the model, ExoIII footprinting, which only detects the specific CF·DNA complex, failed to show significant CF binding to the -27 and -28 mutant promoters (**Fig. 1b**). We conclude that the recognition of base pair identity at the position -28 and to a lesser extend at -27 are important mechanistic elements in defining the CF binding specificity.

The observed binding rate constants in different CF concentration were 0.112–0.180 s^-1^ (*k*_obs,avg_: 0.143±0.023 s^-1^) and 0.281–0.661 s^-1^ (*k*_obs,avg_: 0.375±0.112 s^-1^) for DNA(Cy3,-27) and DNA(Cy3,-28), respectively (**Fig. 3c**; **Fig. S5d,c**; **Table S5**). These values are 3–11-fold larger than the corresponding binding rate of WT DNA(Cy3) (*k*_obs,avg_: 0.036±0.002 s^-1^), but the effect on *k*_obs_ could be because of changes in either the forward (*k*_for_) or reverse (*k*_rev_) reaction rate. To determine *k*_rev_, we mixed the preformed CF·DNA(Cy3,-27) and CF·DNA(Cy3,-28) complexes with unlabeled WT competitor DNA (**Fig. 3d**). Median CF dissociation rates (*k*_rev_) were found as 0.0358±0.0002 s^-1^ [DNA(Cy3,-27)] and 0.3107±0.0056 s^-1^ [DNA(Cy3,-28)], which are significantly larger than from WT DNA(Cy3) (0.0076±0.0002s^-1^) (**Table S5**). Next, *k*_for_ values were calculated (from *k*_obs_=*k*_for_+*k*_rev_) to be 0.107±0.023 s^-1^ [DNA(Cy3,-27)] and 0.064±0.118 s^-1^ [DNA(Cy3,-28)], again somewhat larger than 0.0284±0.0022 s^-1^ for WT DNA(Cy3). The *k*_rev_ values indicate that CF interaction with the base pair -28 is essential for the stability of the specific CF·DNA complex; the substitution of -28 bp caused rapid dissociation of CF from this binding site (*k*_rev_ increased 40-fold vs. wild-type) as well as unfavorable equilibrium constant (*K*_2_=*k*_for_/*k*_rev_=0.21±0.38 vs. WT 3.75±0.37) for the isomerization of initial CF·DNA complex to the final complex, i.e., to high PIFE state (**Table S5**). The substitution of base pair -27 also destabilized the specific CF·DNA complex in the terms of the dissociation rate as *k*_rev_ increased by 4.7-fold. However because the substitution also increased *k*_for_ by 3.8-fold, the equilibrium constants *K*_2_ (3.00±0.66) was little affected. How then the observed PIFE on DNA(Cy3,-27) scaffold was about 3-fold smaller that on the WT DNA(Cy3) (**Fig. 3b**)? One possibility is that the mutation at -27 causes the re-orientation of the CF binding conformation and, thus, only a small PIFE effect.

### Promoter determinants in transcription start site selection

As discussed above, TSS did not change as a response to the 1 bp insertion (between -13/-12) or deletion (−13) at the linker region or rDNA promoter, suggesting that in addition to CF also Pol I contributes to the recognition of the correct TSS. To study further which promoter properties Pol I utilizes to define TSS, we conducted transcription initiation analysis using fork junction, pre-melted and negatively supercoiled rDNA promoters (**Fig. 6a**). We prepared fork promoter FPt1 by annealing ntDNA strand -6/+30 with tDNA strand -15/+30 to address whether Pol I can recognize the sequence of template DNA near the natural TSS. The presence of single stranded template DNA in FPt2 supported Pol I activity (**Fig. 6b**), unlike the fully complementary -15/+30 promoter (**Fig. 4a**). However, Pol I initiates transcription from multiple TSSs (mTSS), the predominant initiation site being near the upstream end of FPt1 (**Fig. 6c**, **e**). Pol I appears thus not to recognize the identity of template DNA nucleotides as part of the TSS recognition mechanism. Although CF was not essential for the transcription activity on FPt1, it stimulated the RNA production rate by 2.7-fold (**Fig. 6c**). Because FPt1 lacks the specific CF binding site, it appears that CF, relying on its unspecific DNA binding mode, attracts Pol I to the vicinity of FPt1 thereby facilitating transcription activity. The presence of single stranded template DNA appears an essential for unspecific Pol I activity because the constructs with extended single stranded ntDNA, specifically FPnt1 and FPnt2, were inactive (**Fig. 6b, e**). Corollary, we conclude that Pol I does not identify template DNA nucleotides.

**Figure 6.**
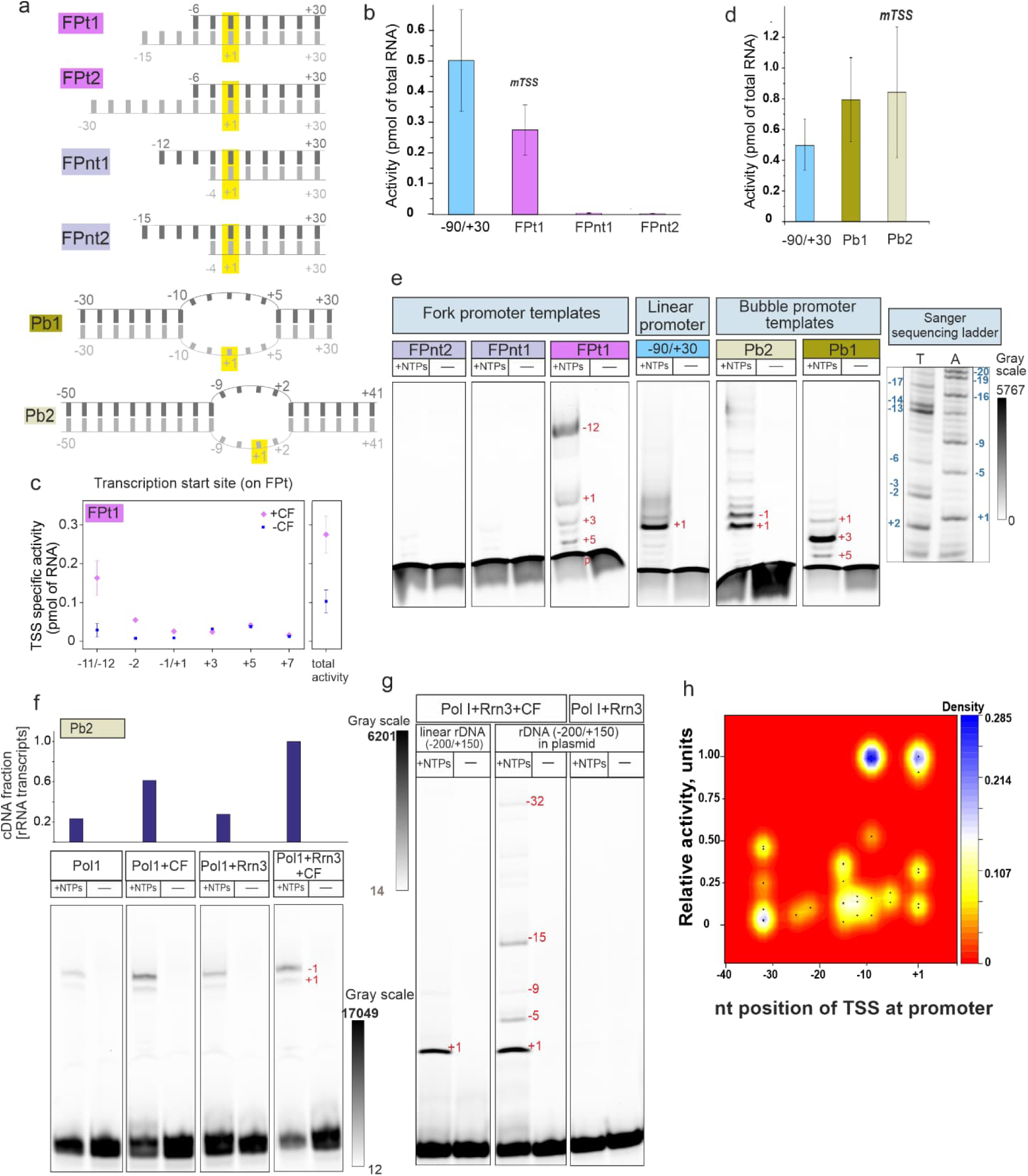
Promoter topology affects the TSS recognition by Pol1. (a) Cartoon represents the structure of fork junction promoters (FPt1, 9 nt overhang in tDNA; FPt2, 24 nt overhang in tDNA; FPnt1, 8 nt overhang in ntDNA; FPnt2, 11 nt overhang in ntDNA) and two pre-melted bubble promoters (Pb1, ntDNA·tDNA mismatch from -10 to +5; Pb2, mismatch from -9 to +2) used to investigate the TSS selection mechanism of Pol I. Numbers indicate the span of each DNA strand, the positions of mismatch regions, and the position of main transcription start site (+1) of WT rDNA promoter. ntDNA and tDNA are shown in dark and light grey, respectively. (b) The transcription initiation activity of fork promoters was measured in the presence of Pol I, Rrn3 and CF. When multiple TSS were observed, as in the case of FPt, Pb1 and Pb2, RNA products were separately quantitated and summed up to obtain the reported total Pol I activity. WT linear rDNA promoter (span -90/+30) was used as a reference control. Reported activities are averages and SD of three independent experiments. (c) Transcription activity on FPt1 was measured with and without CF in the presence of Pol I and Rrn3. Total activity and TSS-specific transcription are shown. (d) Total transcription activity of Pb1 and Pb2 (pre-melted promoters) was measured under the same conditions as in panel b. Data represent averages±SE from three independent experiments. (e) Example PAGE gel visualizes TSS position(s) on different types of promoters. T and A indicate Sanger sequencing ladders prepared using TTP and ATP analogues as chain terminators, respectively. The inferred TSS corresponding to each gel band and the free primer-Cy5.5 are indicated in red font with numbers and p, respectively. (f) Primer extension gel and its quantification (relative cDNA content) of transcription on Pb2 promoter with different components of the Pol I basal system. Transcription activity of Pol I·Rrn3·CF is set to 1; other samples are shown as relative fractions. (g) Example of PAGE gel compares the TSS selection by Pol I on the full-length rDNA promoter (span -200/+150) embedded in a linear or a negatively supercoiled DNA (plasmid). Transcription reaction were performed using basal Pol1 system (Pol I, Rrn3, CF) or holoenzyme (Pol1, Rrn3). (h) Kernel density estimation plot of transcription levels by Pol1·Rrn3·CF complex at each TSS position on a negatively supercoiled promoter (n=8). The y-axis shows relative transcription activity, the x-axis displays TSS positions, and the z-axis (color gradient) indicates TSS density. The calibration bar represents TSS density (probability) at heatmap from 0 to 0.285, with maximum density 1.The presented PAGE gel examples (in panels e, f and g) illustrate the series of transcription reactions with (+NTPs) or without NTPs (−) performed in parallel and run on the same gel.

Next, we assessed TSS on pre-melted promoter scaffolds, which have been used in recent cryo-EM studies to obtain the structure of Pol I·promoter open complex^11,50^. The minimal rDNA promoter, spanning region -30/+30 with a non-template strand mismatch from -10 to +5 (named as Pb1), was active (**Fig. 6d**) and displayed multiple TSS’s within the pre-melted region, most initiation events starting from +3 and less frequent events from +1, +4, or +5 (**Fig. 6e**). A minor translocation of the pre-melted region to -9/+2 (Pb2 promoter) caused a shift in the TSS pattern, the initiation now taking place almost equally efficiently from the positions -1 and +1 (**Fig. 6e**). These finding support the conclusion that Pol I does not possess base identifier function. Presuming that CF interaction with the Pol I and the location of upstream promoter DNA remains normal on the bubble promoters, as is supported by cryo-EM models,^11^ the shift of TSS towards the downstream suggest that Pol I can accommodate differently sized transcription bubbles. The main TSS determining factor on the bubble promoters is the interaction of downstream DNA duplex with Pol I, which guides the first two initiating template bases, near the edge of the duplex, to the active site to provide the template for RNA synthesis. Careful interpretation of structural data, obtained using mismatch promoters, is necessary because these promoters can initiate from non-native TSS. Noteworthy, the transcription rates of pre-melted scaffolds were similar or slightly exceeded that of the native promoter suggesting that the rate-limiting step of the transcription initiation is not the transcription bubble formation (**Fig. 6d**). The activity on Pb2 was stimulated by CF but not absolutely dependent on its presence (**Fig. 6f**).

We finally determined TSS of a full-length rDNA promoter (−200/+150) embedded to a plasmid. Data shows that the Pol I basal transcription machinery exhibits mTSSs on the plasmid template (**Fig. 6g**, **h**). The TSS distribution fluctuate along the promoter between experimental repeats (n=8), probably reflecting variable levels of negative supercoiling in different plasmid promoter preparations. The highest frequency of TSS was found at promoter positions +1, -9 and -15. In contrast, the linear template consistently initiated from +1 (**Fig. 5b**). Initiation on the plasmid and linear promoter is strictly dependent on the presence of CF. Negative supercoiling stimulates processes that require DNA helix opening, such as transcription and replication initiation. The relaxed TSS selectivity on the plasmid template could be rationalized by assuming that the inherently loose interaction/connection between CF and Pol I allows Pol I to land to the DNA on different positions in the vicinity of CF. The double-stranded DNA can then enter to the cleft of Pol I leading to at least temporal DNA strand separation and transcription bubble formation, and hence RNA synthesis. On non-supercoiled DNA the melting of DNA requires more subtle and stable positioning of Pol on the promoter to allow the transcription bubble formation and initiation of RNA synthesis.

### Structural dynamics of Pol I closed complex

We next probed the structural dynamics in Pol I closed complex, to understand better the reaction path to the open complex, by performing molecular dynamic simulations of the system. The simulations were started from one of the two closed complex conformations (CC2) reconstructed by Sadian et al. from cryo-EM data.^11^ We selected CC2 over CC1 because of its better resolution (avg. 2.9 vs 3.8 Å). One essential feature of CC has been suggested to be the open conformation of the clamp/cleft in Pol I ^11^. This conformation is believed to be connected with the stable anchoring of DNA-mimicking loop (DML)/expander of A190 subunit and the C-terminal domain of A12.2 (CTD A12.2) inside the DNA-binding cleft of Pol I, as well as the partial unfolding of the bridge helix (in A190). Upon the formation of the OC, clamp/cleft adopts a more narrow closed conformation, the DML/expander is expelled from the DNA-binding cleft and the bridge helix (BH) refolds to more helical structure. The subsequent initiation of RNA synthesis also mobilizes the CTD A12.2 as indicating by the loss of its density in cryo-EM data ^11^. We first analyzed the MD trajectories for the behavior of these domains and conformations. The visual and distance (e.g., from the expander to reference points in Pol I as depicted in **Fig. S6a-c**) based analysis of the trajectories indicated that both the DML/expander and CTD A12.2 remained relatively immobile in the DNA-binding cleft. The observation is not surprising because the simulation time (3×80 ns) was relatively short to observe larger domain movements and binding or unbinding events. However, the inspection of clamp/cleft suggested significant dynamical changes. Specifically, the cleft width (measured between C-alpha atoms of residue Thr429 in A190 and residue Asn423 in A135 as previously in Sadian et al.^11^) varied between 31 Å and 38 Å (**Fig. S6a-c**), thus, approaching the values measured in the cryo-EM models for the closed clamp state (31.6 Å in the PDB model 6rwe depicting OC2) and open clamp state [39.5 Å in the PDB models 6rqh (CC1) and 6rql (CC2)], respectively. MD data thus suggests that the presence of the DML/expander and CTD A12.2 in the DNA-binding cleft does not restrict the clamp dynamics. We then inspected the BH and found the expansion of the unfolded region in the middle of the BH in the simulations 2 and 3 (**Fig. S7a**). Based on the positive Pearson correlation coefficient (0.45, p-value 5.8E-09), there appears to be a moderately strong correlation between the unfolding of the BH and the opening of the clamp, and *vice versa*.

CF interacts with Pol I-bound Rrn3 through the Zn-ribbon of Rrn7. In the MD simulations, the position of the Rrn7 Zn-ribbon relative to Rrn3 and the surrounding Pol I structure remained stable (**Fig. S7b-d**). Additionally, the acidic loop of Rrn3 consistently remained wrapped around the Rrn7 Zn-ribbon, thus providing consistent stabilization for the main link of Pol I·Rrn3·CF complex.

Pol I CCs bends the promoter significantly at base pairs -11 to -7, the distortion of the double helix potentially driving subsequent DNA melting and OC formation.^11^ In MD trajectories, the bending angle of the promoter stayed mostly between 120° and 160° (180° is straight promoter), indicating that the binding of the downstream DNA between the clamp and lobe domains of Pol I, the binding of the upstream DNA to CF and the relative positioning Pol I and CF define relatively stable constrain to the geometry of the promoter in the CC state (**Fig. S8a,b**). The base pairing within the kinked region was indeed unstable, especially the template DNA base -8 showing wide-ranging mobility between the canonical base pairing (**Fig. S8c**) and flipping out of the base stack (**Fig. S8d**). In the fully flipped out state, the template DNA base -8 interacts with Arg448 and Met451 in the A135 protrusion domain. MD trajectories thus align with the mechanism that, as part of the CC-to-OC transition, DNA bases within the kinked promoter region (−11 to -7) flip out from the double helix and are stabilized by interactions with the surrounding protein, until the expanding region of the single-stranded template DNA leads to its loading into the DNA-binding cleft.

### Interactions of the promoter and CF in MD simulations

The N-terminal DNA-binding helical bundle (DBHB) of Rrn11 and the loop between helices α8 and α9 contact DNA in the major and minor groove. In the simulations, Rrn11 mostly interacts with the phosphate backbone of the promoter DNA and the interactions with the DNA remain rather similar to those observed in the cryo-EM structure. Sadian et al. ^11^ identified only one residue that makes base specific contacts; Arg11 which interacts with two adjacent adenines in positions -22 and -21. In simulations 1 and 2, Arg11 remains interacting with these adenines, often to the backbone phosphates, but in simulation 3 it rotates away from the adenines and does not participate in the interaction with the promoter DNA (**Fig. 7a,b**).

**Figure 7.**
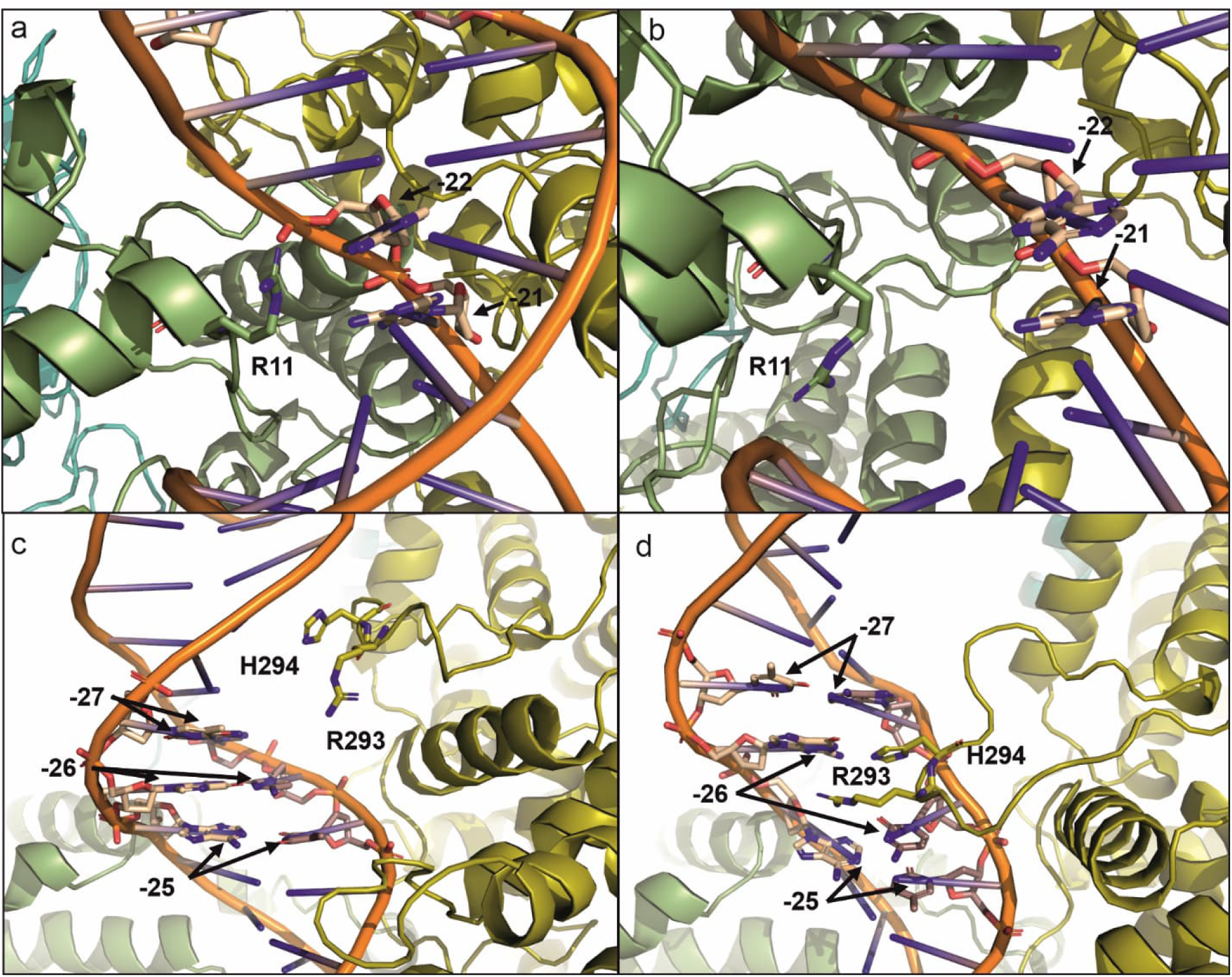
Snapshots from MD simulation trajectory detailing the interaction of CF with rDNA promoter. (a) The interaction of Rrn11 residue Arg11 (R11) with the promoter bp -22 and bp -21 in simulation 1. (b) Rrn11 residue Arg11 (R11) rotated away from bp -22 and bp -21 in simulation 3. (c) Typical orientation of the loop between α7 and α8 of Rrn7 from simulation 1.The loop retracted away from the DNA major groove. (d) The insertion of Rrn7 residues Arg293 and Arg294 between the DNA bp -27, -26 and -25 as observed in simulation 3. The loop between α7 and α8 of Rrn7 inserted into the DNA major groove.

The Rrn7 C-terminal cyclin fold contacts the upstream DNA with the loop between α7 and α8, including the insertion of R293 between the DNA bases at positions -27 and -26.^11^ These and DBHB of Rrn11 interactions with promoter DNA bases and backbone apparently impose significant widening of the major groove between positions -27 and -22. Collectively CF and Pol I cause bending and distortion of the promoter DNA within in vicinity the CF binding site. We anticipate that these DNA distortions are energetically costly and require sophisticated network of interactions to be compensated by binding energy. As an indirect evidence in favor of this hypothesis, we observed that the loop between α7 and α8 of Rrn7 retracted away from the major groove in most simulations, potentially suggesting that the force field thereby evaded the adverse geometrical/clash energies (**Fig. 7c**). The notable exceptions was WT simulation 3, in which the loop, after initial retraction, re-inserted residues Arg293 and His294 in between of the DNA bp -27, -26 and -25 (**Fig. 7d**). The loop insertion apparently also caused substantial the DNA major groove between positions -27 and -22 (**Fig. S9a,b**). Notably, this is the same simulation where the interaction of Rrn11 Arg11 with DNA near bp -22/-21 was lost.

In an attempt to probe the structural basis of -28 and -27 promoter mutations on the complex formation with CF, we performed introduced the corresponding substitutions to the DNA and repeated the MD simulations. The base pairs -28 and -27 interacted only with Rrn11 Lys182 in the simulations. The interaction was rather infrequent, taking place in 15–30% of total trajectory length, and often water mediated (at least 50% of all cases). Thus, the -28 and -27 mutations appear not to affect the promoter interaction with Rrn11. The interaction of Rrn7 with base pairs -28 and -27 was more extensive. In the WT promoter simulations, the base pair at -28 interacts with residues Arg293, His294, Thr295 and Arg297. In the WT simulation 3, in which Arg293 and His294 insert themselves in-between the base pairs -27, -26 and -25, the interaction with base pair -28 is lost. This corresponds to the overall lower interaction frequency of the WT promoter at base pair -28 with Rrn7. When the base pair at -28 is mutated, its interaction frequency with Arg293, His294, Thr295 and Arg297 is increased, again probably because these simulations did not show the re-insertion of the loop between Rrn7 α7 and α8 deep into the promoter major groove and, specifically, the penetration of Arg293/His294 between the base pairs -27, -26 and -25 (**Fig. S9c**). In WT promoter, the base pair at -27 interacts with Rrn7 residues Asn222, Arg293, His294, Thr295 and Gly296. When the base pair -27 is mutated, the most significant change is the increased interaction frequency with Arg293. As expected, when Arg293 and His294 inserted between the bases -27, -26 and -25 in the WT simulation 3, the calculated interaction frequency between the base pair -27 and Arg293/His294 increased. In contrast, the mutated base pair -27 does not interact significantly with His294 and the CF loop remains outside the DNA major groove (**Fig. S9d**). Overall, the mutation of base pairs -27 and -28 affected multiple ways the simulated interaction modalities of the promoter DNA and Rrn7. We speculate that the insertion (or retraction in the reverse reaction) of Rrn7 loop between α7 and α8 to the major groove of the promoter (between positions -28 and -21) may be the equivalent of the second step (isomerization step) in the two-step binding mechanism of CF to the promoter. This insertion is not required for the initial binding of CF to the promoter but is driving the final maturation/isomerization of the promoter·CF complex. Within this framework, consistent with our SF data, the mutated -28 and -27 base pairs have small (if any) effect on the initial CF binding but decrease the stability of the loop insertion in the promoter major groove.

## DISCUSSION

In this study, we provide novel functional and kinetic data on Pol I transcription initiation, offering insights into promoter recognition and transcription start site selection by CF and Pol I. These findings significantly enhance our understanding of Pol I initiation, with the kinetic parameters providing a key advance in elucidating the dynamics of this process.

CF recognizes the rDNA promoter and recruits Pol I on it. To provide further insight on how CF recognizes its binding site, we employed fluorescence-based PIFE assay to perform, for the first time, the detailed kinetic analysis of the CF·promoter complex formation and stability. We estimated the apparent *K*_D_ value for the interaction to be about 50–100 nM. This value deviates significantly from a previously reported *K*_D_ value of 0.4 pM, obtained using EMSA technique ^34^. We suggest that the difference in the estimates is explained by the “caging” effect of the gel matrix that is known to stabilize protein·DNA complexes ^51^. Our PIFE data further indicated that CF dissociates moderately slowly from the promoter (median reaction time 93.6±2.1 s, parameter *k*_rev_ in **Table S5**), explaining why ExoIII treatment produced CF footprint on the promoter (this study and ^11^). The determined CF dissociation rate is also consistent with promoter switch (or promoter commitment) experiments demonstrating, within 30 min timescale, the “unstable” and “stable” binding of CF and Pol I to the promoter in the absence and presence of UAF (and TBP), respectively^18^. Consistently, the lack of exoIII footprint for Pol I in our experiments indicate unstable CF mediated recruitment of Pol I to the promoter. In the context of CF·promoter kinetic parameters, we propose that the documented activating effects of UAF on the Pol I apparatus recruitment and activity ^18,52^, at least in large part, arise from the stabilization of CF·promoter complex, more likely via decreased rate of CF dissociation.

The lack of sequence conservation and computationally predicted physical properties of rDNA promoters culminated to a hypothesis that CF in yeast and its functional analogues in other species, including SL1/TIF-1B in humans, mouse and *Acanthmoeba castellanii*, recognize conserved structural anomalies, such as curvature, bendability and (minor) groove width, in the rDNA promoter ^34,53^. A recent high-resolution cryo-EM study indicated that CF in fact contacts two widened DNA grooves, specifically the major groove from position -28 to -21 and the adjacent minor groove from -24 to -19, in the promoter that is additionally bent by about 30 degrees at around position -20 ^11^. However, it remained unclear whether these structural perturbations pre-exist in the free rDNA promoter or are they induced by CF binding? The binding rate analysis can be diagnostic for the mechanism. Three scenarios can be considered to explain the connection between CF binding and the promoter structure. In the first scenario, CF binds directly to a pre-existing perturbed promoter, for simplicity assumed to be the bent promoter. In the second, the free promoter exists in equilibrium between straight and bent conformations, with CF preferentially binding to the bent form. In the third scenario, the free promoter is straight, and the protein binds to it, inducing a conformational change in the initial protein·promoter complex that leads to promoter bending. Our data exclude the first scenario, as it predicts that the binding rate would increase with increasing CF concentration. However, we observed a constant binding rate across all concentrations tested. The second scenario suggests a biphasic binding process: in the rapid phase, the protein binds to the bent promoters present in the equilibrium with the straight form. This would then shift the equilibrium toward the formation of additional bent promoters, which are bound by CF during the slower phase of the reaction. Yet, our experiments revealed only a single-phase binding reactions. The third scenario, basically the induced-fit binding, predicts a single-phase reaction, with the binding rate remaining uniform across different protein concentrations, matching our experimental observations.

Several aspects of our data suggest that the induced-fit mechanism of CF involves, at least in part, events within the DNA major groove (from -28 to -21) of the promoter, including groove deformation and the recognition of specific bases. Specifically, we observed that the truncation of the promoter upstream to -28 elevated both the CF binding affinity (**Fig. 4d**) and the transcription initiation activity (**Fig. 4a**), presumably by facilitating the major groove widening by CF. The substitution of the major groove base pairs -28 and -27 impaired CF·promoter complex as evident by the increased dissociation rate (**Fig. 3d**), consistent with the importance of specific groove interactions for the stability of the complex. Because these substitutions did not decrease the binding rate, the specific interactions with -28 and -27 are less important for the CF binding step, i.e., initial complex formation. MD simulations supported the experimental findings by revealing dynamic interactions, including the binding and unbinding of the Arg293/His294-harbouring loop (between α7 and α8 in Rrn7) within the DNA major groove. In contrast, the interaction of CF with the adjacent minor groove (from -24 to -19) was more stable in the MD simulations. Previous study found that simultaneous substitutions of two to four residues within the minor groove impair CF·promoter complex, thus highlighting the importance of base recognition also in the minor groove ^11^. Overall, the induced fit mechanism here proposed unifies the deformability and base-specific recognition hypotheses for CF specificity: the initial binding of CF triggers structural deformation of the promoter DNA, especially within the DNA major groove, allowing CF to establish specific stabilizing interactions with the nearby bases. The fact that base-recognizing residues in CF, especially N209, R293 and H294 in Rrn7 ^11^, locate to flexible loops may be an essential element of the induced-fit mechanism of CF, allowing the penetration of protein residues into the DNA grooves. We also note that the placement of base-recognition elements to the protein loops has been indicated to facilitate the rapid co-evolution of transcription factors and their promoter binding sites ^54^, the phenomenon potentially contributing to the apparent lack of inter-species conservation of rDNA promoter sequences.

After the binding of CF to the rDNA promoter (specificity box), CF recruits Pol I·Rrn3 complex next to it ^9,10,49,55^. However, the recruitment in our experimental conditions appears transient as no ExoIII footprint for Pol I was observed (**Fig. S3b**,**c**). Structural studies have not identified any specific promoter bases that are selectively recognized by Pol I as a part of the initiation mechanism ^10–12,49,50^. However, the randomization of the promoter sequence at the Pol I binding site impaired transcription activity suggesting that Pol I cannot initiate on any sequence ^9,56,57^. We sought to dissect further the potential base-specificity of Pol I by studying transcription initiation efficiency and transcription start site (TSS) selection using bubble (non-template DNa mismatch) and fork promoter as experimental models that have regions of exposed single stranded DNA (**Fig. 6**). We found no evidence that Pol I would recognize specific bases, and instead the TSS was determined by the topology of the promoter model, i.e., the location of the single-stranded template DNA and downstream duplex. The promoter DNA is sharply kinked ∼60° between the positions -11 and -7 in the Pol I closed complex to avoid clashing of the downstream DNA with the clamp domain of Pol I; the kink may also provide essential destabilization of DNA duplex at the onset of transcription bubble melting ^9–11,49^. Indeed, our MD simulations showed unstable base-pairing at the kinked promoter region (**Fig. S8c**,**d**). When we moved the CF binding site 1 bp closer or further from the TSS (+1) in the promoter, the initiation activity decreased by about 85 % and 65 %, respectively, but the TSS was not changed. Previously, the dislodging of the CF binding site by ±5 bp caused complete elimination of the activity as did also the full substitution of the sequence from -11 to -7^11^. The correct distance between the strong CF binding site and the kinked sequence (from -11 to - 7) is thus one critical element of functional rDNA promoter. The current working hypothesis is that the sequence from -11 to -7 needs to be ‘bendable’ to allow the formation of the promoter kink in CC, the initiation of DNA strand separation (melting) and the loading of the template DNA strand into the active site cleft of Pol I ^11^. However, integrated structural, computational and functional studies will be needed to dissect the exact structure-function relationship within this promoter region. The initiation of the DNA melting is followed the bubble expansion to the downstream direction ^58^. When enough DNA melting and template strand loading has occurred to allow sufficiently stable docking of the templating nucleobases to the Pol I active center, the first two NTPs bind to the active site and RNA synthesis initiates. Considering that we did not find Pol I - dependent base recognition and that mutant rDNA promoter containing deletion from the position -5 to +128 was reported partially active ^57^, the bubble expansion process and the selection of the exact TSS appears to be more defined by the promoter DNA geometry, especially distance from the bubble initiation site (near positions -11 and -7), than the specific sequence near the TSS.

In conclusion, CF primes the transcription of rDNA by induced-fit mechanism based recognition of its sequence-specific binding site on the promoter and the subsequent recruitment of Pol I·Rrn3 complex. Pol I machinery in addition to the specific binding site for the CF responds to DNA conformation. Interestingly, the movement of RNA polymerase along the DNA generates negative supercoils behind RNA polymerase and positive supercoils in front, as described by the twin-supercoiled-domain model ^58,59^. This model predicts that two domains of DNA supercoiling are generated during transcription elongation, provided that the RNA polymerase meets resistance to rotate freely around the template, and that DNA rotation is hindered ^60^. Transcription-generated supercoils can, in turn, enhance or impede the transcriptional process: negative supercoils facilitate transcription initiation by enabling promoter melting and enhancing the binding of regulatory factors, whereas positive supercoils aid the elongation of RNA polymerases by destabilizing DNA-bound proteins. However, excessive negative or positive supercoils can also repress transcription. We showed that changes in DNA topology led to selection of alternative TSSs at native promoter ^61,62^. Our *in vitro* transcription data with full length (−200/+150) promoter inserted in plasmid constantly demonstrated variable multiple TSSs while the linear prototype of promoter showed single TSS at +1 (**Fig. 6g**,**h**). Furthermore, transcription initiation from circular (negatively supercoiled) DNA template was exclusively CF-dependent and promoter-specific that aligns with our results collected at linear promoter DNA. This, in turn, indicates the major role of physical properties of DNA (torsional forces etc.) in selective transcription initiation on rDNA promoter by Pol I. Next, pre-formed mismatched regions (pre-melted promoters), resulted in TSS transition to loci different from +1 on the promoter, or even initiation from multiple sites. Thus, our findings confirm that rDNA promoter properties and sequence conservation in the core element region define efficient rDNA transcription, including transcription rate, specificity and TSS selection. Referring to previously published works (reviewed in ^55,63–68)^ and our data, we conclude that Pol I machinery in addition to the specific binding site for the CF relies on a combination of physical features of DNA, the “bendability” of upstream promoter DNA and the ‘‘meltability’’ of the region around the TSS.

## Acknowledgement

We acknowledge Turku Protein Core (TuProtCore) for providing access to mass photometer and stopped flow instruments. Fluorescence scanning of the gels was performed at the Cell Imaging and Cytometry Core, Turku Bioscience Centre, Turku, Finland, with the support of Biocenter Finland. The authors also wish to acknowledge CSC – IT Center for Science, Finland, for computational resources.

## Data availability

Molecular dynamic simulations data is available from Zenodo.org (EU Open Research Repository) using the DOI 10.5281/zenodo.14001801.

## Supplementary materials

Supplementary Tables 1–5

Supplementary Figures 1–9

During the preparation of this work the authors used ChatGPT in order to improve language and readability. After using this tool, the authors reviewed and edited the text and take full responsibility for the content of the publication.

## REFERENCES

1. Ferreira, R., Schneekloth, J. S., Panov, K. I., Hannan, K. M. & Hannan, R. D. Targeting the RNA polymerase i transcription for cancer therapy comes of age. Cells 9 (2):266 10.3390/cells9020266 (2020).

2. Warner, J. R. The economics of ribosome biosynthesis in yeast. Trends in Biochemical Sciences vol. 24 10.1016/S0968-0004(99)01460-7 (1999).

3. Mayer, C., Zhao, J., Yuan, X. & Grummt, I. mTOR-dependent activation of the transcription factor TIF-IA links rRNA synthesis to nutrient availability. Genes Dev 18, (2004).

4. Farley-Barnes, K. I., Ogawa, L. M. & Baserga, S. J. Ribosomopathies: Old Concepts, New Controversies. Trends in Genetics 35, 10.1016/j.tig.2019.07.004 (2019).

5. Bywater, M. J. et al. Inhibition of RNA Polymerase I as a Therapeutic Strategy to Promote Cancer-Specific Activation of p53. Cancer Cell 22, (2012).

6. Pelletier, J., Thomas, G. & Volarević, S. Ribosome biogenesis in cancer: new players and therapeutic avenues. Nat Rev Cancer 18, 51–63 (2018).

7. Rossetti, S., Wierzbicki, A. J. & Sacchi, N. Cell Cycle Mammary epithelial morphogenesis and early breast cancer. Evidence of involvement of basal components of the RNA Polymerase I transcription machinery. (2016) doi:10.1080/15384101.2016.1215385.

8. Pilsl, M. & Engel, C. Structural basis of RNA polymerase I pre-initiation complex formation and promoter melting. Nature Communications 2020 11:1 11, 1–10 (2020).

9. Engel, C. et al. Structural Basis of RNA Polymerase I Transcription Initiation. Cell 169, (2017).

10. Sadian, Y. et al. Structural insights into transcription initiation by yeast RNA polymerase I. EMBO J 36, (2017).

11. Sadian, Y. et al. Molecular insight into RNA polymerase I promoter recognition and promoter melting. Nature Communications 2019 10:1 10, 1–13 (2019).

12. Tafur, L. et al. The cryo-EM structure of a 12-subunit variant of RNA polymerase I reveals dissociation of the A49-A34.5 heterodimer and rearrangement of subunit A12.2. Elife 8, (2019).

13. Hannig, K. et al. The C-terminal region of Net1 is an activator of RNA polymerase I transcription with conserved features from yeast to human. PLoS Genet 15, (2019).

14. Shou, W. et al. Net1 stimulates RNA polymerase I transcription and regulates nucleolar structure independently of controlling mitotic exit. Mol Cell 8, (2001).

15. Hernandez, N. TBP, a universal eukaryotic transcription factor? Genes and Development vol. 7 Preprint at 10.1101/gad.7.7b.1291 (1993).

16. Kwan, J. Z. J., Nguyen, T. F. & Teves, S. S. TBP facilitates RNA Polymerase I transcription following mitosis. RNA Biol 21, 42–51 (2024).

17. Gorski, J. J. et al. A novel TBP-associated factor of SL1 functions in RNA polymerase I transcription. EMBO J 26, 1560–1568 (2007).

18. Steffan, J. S., Keys, D. A., Dodd, J. A. & Nomura, M. The role of TBP in rDNA transcription by RNA polymerase I in Saccharomyces cerevisiae: TBP is required for upstream activation factor-dependent recruitment of core factor. Genes Dev 10, 2551–2563 (1996).

19. Siddiqi, I., Keener, J., Vu, L. & Nomura, M. Role of TATA Binding Protein (TBP) in Yeast Ribosomal DNA Transcription by RNA Polymerase I: Defects in the Dual Functions of Transcription Factor UAF Cannot Be Suppressed by TBP. Mol Cell Biol 21, 2292–2297 (2001).

20. Bric, A., Radebaugh, C. A. & Paule, M. R. Photocross-linking of the RNA polymerase I preinitiation and immediate postinitiation complexes. Implications for promoter recruitment. Journal of Biological Chemistry 279, (2004).

21. Al-Khouri, A. M. & Paule, M. R. A Novel RNA Polymerase I Transcription Initiation Factor, TIF-IE, Commits rRNA Genes by Interaction with TIF-IB, Not by DNA Binding. Mol Cell Biol 22, 750–761 (2002).

22. Radebaugh, C. A. et al. TATA box-binding protein (TBP) is a constituent of the polymerase I-specific transcription initiation factor TIF-IB (SL1) bound to the rRNA promoter and shows differential sensitivity to TBP-directed reagents in polymerase I, II, and III transcription factors. Mol Cell Biol 14, 597–605 (1994).

23. Steffan, J. S., Keys, D. A., Dodd, J. A. & Nomura, M. The role of TBP in rDNA transcription by RNA polymerase I in Saccharomyces cerevisiae: TBP is required for upstream activation factor-dependent recruitment of core factor. Genes Dev 10, 2551–2563 (1996).

24. Keys, D. A. et al. RRN6 and RRN7 encode subunits of a multiprotein complex essential for the initiation of rDNA transcription by RNA polymerase I in Saccharomyces cerevisiae. Genes Dev 8, 2349–2362 (1994).

25. Lalo, D., Steffan, J. S., Dodd, J. A. & Nomura, M. RRN11 encodes the third subunit of the complex containing Rrn6p and Rrn7p that is essential for the initiation of rDNA transcription by yeast RNA polymerase I. J Biol Chem 271, 21062–21067 (1996).

26. Steffan, J. S., Keys, D. A., Vu, L. & Nomura, M. Interaction of TATA-Binding Protein with Upstream Activation Factor Is Required for Activated Transcription of Ribosomal DNA by RNA Polymerase I in Saccharomyces cerevisiae In Vivo . Mol Cell Biol 18, (1998).

27. Yamamoto, R. T., Nogi, Y., Dodd, J. A. & Nomura, M. RRN3 gene of Saccharomyces cerevisiae encodes an essential RNA polymerase I transcription factor which interacts with the polymerase independently of DNA template. EMBO Journal 15, (1996).

28. Keener, J., Josaitis, C. A., Dodd, J. A. & Nomura, M. Reconstitution of Yeast RNA Polymerase I Transcription in Vitro from Purified Components. Journal of Biological Chemistry 273, 33795–33802 (1998).

29. Bedwell, G. J., Appling, F. D., Anderson, S. J. & Schneider, D. A. Efficient transcription by RNA polymerase I using recombinant core factor. Gene 492, 94–99 (2012).

30. Keener, J., Josaitis, C. A., Dodd, J. A. & Nomura, M. Reconstitution of yeast RNA polymerase I transcription in vitro from purified components: TATA-binding protein is not required for basal transcription. Journal of Biological Chemistry 273, (1998).

31. Moreno-Morcillo, M. et al. Solving the RNA polymerase I structural puzzle. Acta Crystallogr D Biol Crystallogr 70, 2570–2582 (2014).

32. Rigaut, G. et al. A generic protein purification method for protein complex characterization and proteome exploration. Nat Biotechnol 17, 1030–1032 (1999).

33. Mukai, T. et al. Highly reproductive Escherichia coli cells with no specific assignment to the UAG codon. Sci Rep 5, 9699 (2015).

34. Jackobel, A. J., Zeberl, B. J., Glover, D. M., Fakhouri, A. M. & Knutson, B. A. DNA binding preferences of S. cerevisiae RNA polymerase I Core Factor reveal a preference for the GC-minor groove and a conserved binding mechanism. Biochimica et Biophysica Acta (BBA) - Gene Regulatory Mechanisms 1862, 194408 (2019).

35. Belogurov, G. A. et al. Membrane-Bound Pyrophosphatase of *Thermotoga maritima* Requires Sodium for Activity. Biochemistry 44, 2088–2096 (2005).

36. Carey, M. F., Peterson, C. L. & Smale, S. T. The primer extension assay. Cold Spring Harb Protoc 8, (2013).

37. Keener, J., Josaitis, C. A., Dodd, J. A. & Nomura, M. Reconstitution of Yeast RNA Polymerase I Transcription in Vitro from Purified Components. Journal of Biological Chemistry 273, 33795–33802 (1998).

38. Rueden, C. T. et al. ImageJ2: ImageJ for the next generation of scientific image data. BMC Bioinformatics 18, 529 (2017).

39. Metzger, W. & Heumann, H. Footprinting with exonuclease III. Methods Mol Biol 148, (2001).

40. Young, G. et al. Quantitative mass imaging of single biological macromolecules. Science (1979) 360, (2018).

41. Flomenbom, O. et al. Stretched exponential decay and correlations in the catalytic activity of fluctuating single lipase molecules. Proc Natl Acad Sci U S A 102, (2005).

42. Mäkinen, J. J. et al. The mechanism of the nucleo-sugar selection by multi-subunit RNA polymerases. Nat Commun 12, 796 (2021).

43. Lu, C. et al. OPLS4: Improving Force Field Accuracy on Challenging Regimes of Chemical Space. J Chem Theory Comput 17, 4291–4300 (2021).

44. Hwang, H. & Myong, S. Protein induced fluorescence enhancement (PIFE) for probing protein–nucleic acid interactions. Chem. Soc. Rev. 43, 1221–1229 (2014).

45. Ha, J. H., Capp, M. W., Hohenwalter, M. D., Baskerville, M. & Record, M. T. Thermodynamic stoichiometries of participation of water, cations and anions in specific and non-specific binding of lac repressor to DNA. Possible thermodynamic origins of the ‘glutamate effect’ on protein-DNA interactions. J Mol Biol 228, (1992).

46. Purich, D. L. Enzyme Kinetics: Catalysis & Control. Enzyme Kinetics: Catalysis & Control (2010). doi:10.1016/C2009-0-61154-5.

47. Ross, W. & Gourse, R.L. Sequence-independent upstream DNA-alphaCTD interactions strongly stimulate *Escherichia coli* RNA polymerase-*lacUV5* promoter association. Proc. Natl. Acad. Sci. U. S. A., 102, 291–296 (2005).

48. Davis, C.A., Capp, M.W., Record, M.T. & Saecker, R.M. The effects of upstream DNA on open complex formation by Escherichia coli RNA polymerase. Proc. Natl. Acad. Sci. U. S. A., 102, 285–290 (2005).

49. Han, Y. et al. Structural mechanism of ATP-independent transcription initiation by RNA polymerase I. Elife 6, (2017).

50. Tafur, L. et al. Molecular Structures of Transcribing RNA Polymerase I. Mol Cell 64, (2016).

51. Fried, M. G. & Liu, G. Molecular sequestration stabilizes CAP–DNA complexes during polyacrylamide gel electrophoresis. Nucleic Acids Res 22, 5054–5059 (1994).

52. Keys, D. A. et al. Multiprotein transcription factor UAF interacts with the upstream element of the yeast RNA polymerase I promoter and forms a stable preinitiation complex. Genes Dev 10, 887–903 (1996).

53. Marilley, M., Radebaugh, C., Geiss, G., Laybourn, P. & Paule, M. DNA structural variation affects complex formation and promoter melting in ribosomal RNA transcription. Molecular Genetics and Genomics 267, 781–791 (2002).

54. Berg, J., Willmann, S. & Lässig, M. Adaptive evolution of transcription factor binding sites. BMC Evol Biol 4, 42 (2004).

55. Roeder, R. G. 50+ years of eukaryotic transcription: an expanding universe of factors and mechanisms. Nature Structural and Molecular Biology vol. 26 783–791 Preprint at 10.1038/s41594-019-0287-x (2019).

56. Choe, S. Y., Schultz, M. C. & Reeder, R. H. *In vitro* definition of the yeast RNA polymerase I promoter. Nucleic Acids Res 20, 279–285 (1992).

57. Kulkens, T., Riggs, D. L., Heck, J. D., Planta, R. J. & Nomura, M. The yeast RNA polymerase I promoter: ribosomal DNA sequences involved in transcription initiation and complex formation in *in vitro*. Nucleic Acids Res 19, 5363–5370 (1991).

58. Kahl, B. F., Li, H. & Paule, M. R. DNA melting and promoter clearance by eukaryotic RNA polymerase I 1 1Edited by R. Ebright. J Mol Biol 299, 75–89 (2000).

59. Janissen, R., Barth, R., Polinder, M., van der Torre, J. & Dekker, C. Single-molecule visualization of twin-supercoiled domains generated during transcription. Nucleic Acids Res 52, 1677–1687 (2024).

60. Tsao, Y.-P., Wu, H.-Y. & Liu, L. F. Transcription-driven supercoiling of DNA: Direct biochemical evidence from in vitro studies. Cell 56, 111–118 (1989).

61. Drolet, M. Growth inhibition mediated by excess negative supercoiling: the interplay between transcription elongation, R-loop formation and DNA topology. Mol Microbiol 59, 723–730 (2006).

62. Chong, S., Chen, C., Ge, H. & Xie, X. S. Mechanism of Transcriptional Bursting in Bacteria. Cell 158, 314–326 (2014).

63. Girbig, M., Misiaszek, A. D. & Müller, C. W. Structural insights into nuclear transcription by eukaryotic DNA-dependent RNA polymerases. Nat Rev Mol Cell Biol 23, 603–622 (2022).

64. Jackobel, A. J., Han, Y., He, Y. & Knutson, B. A. Breaking the mold: structures of the RNA polymerase I transcription complex reveal a new path for initiation. Transcription 9, 255–261 (2018).

65. Khatter, H., Vorländer, M. K. & Müller, C. W. RNA polymerase I and III: similar yet unique. Curr Opin Struct Biol 47, 88–94 (2017).

66. Schwank, K. et al. Features of yeast RNA polymerase I with special consideration of the lobe binding subunits. Biol Chem 404, 979–1002 (2023).

67. Jochem, L., Ramsay, E. P. & Vannini, A. RNA polymerase I, bending the rules? EMBO J 36, 2664–2666 (2017).

68. Pitts, S. & Laiho, M. Regulation of RNA Polymerase I Stability and Function. Cancers (Basel) 14, 5776 (2022).

69. Hossain, K. A. et al. How acidic amino acid residues facilitate DNA target site selection. Proceedings of the National Academy of Sciences 120, (2023).

70. Pettersen, E. F. et al. UCSF Chimera—A visualization system for exploratory research and analysis. J Comput Chem 25, 1605–1612 (2004).

