## Supplemental material for "Determinants of transcription initiation efficiency and start site selection by RNA polymerase I"

Supplementary Tables 1–5
Supplementary Figures 1–9

**Table S1.** ***E.coli*** **protein expression vectors.**

| Name | Description | Resistance |
|---|---|---|
| pAM037 | Expression construct of 6His-TEV-SceRrn3 in modified pET15<br>Rrn3 cloned from INV *S.cerevisiae* yeast in modified pET15 using NcoI + SacI sites | Kan |
| pAM031 | Modified pET15. Expression of Rrn6 subunit of CF (C-terminal TEV-$His_{10}$) | Amp |
| pAM033 | Co-expression of untagged *S.cerevisiae* Rrn7 and Rrn11 in *S.cerevisiae* in modified pET36b | Kan |
| pAM036 | *S.cerevisiae* ribosomal promoter fragment -200/+150 in pUC18 using SacI+HindIII. | Amp |
| pAM041 | *S.cerevisiae* non-promoter fragment (from ATG13 gene) in pUC18 using SacI+HindIII. | Amp |

**Table S2. DNA oligonucleotides used for primer extension assay and PCR-based preparation of transcription templates.**

| Name | Sequence Sequence from 5' to 3' | Direction | Label | Primer works with |
|---|---|---|---|---|
| OP03 | [Atto680]TCGAATTCGTTTCCAAACTC | Reverse | 5'-Atto680 | Primer extension (downstream TSS) |
| OP018 | [Cy5.5]CGAACTTGTCTTCAACTGC | Reverse | 5' Cyanine5.5 [Cy5.5] | Primer extension, Sanger sequencing (downstream TSS) |
| OP21 | [Cy5]TGGAGAATAGCTTAAATTGAAG | Reverse | 5' Cyanine [Cy5] | Primer extension (upstream TSS) |
| EAS13 | CGGCTCGTATGTTGTGTG | Forward | — | pAM036 and PAM041 to amplify DNA template |
| EAS14 | TTAAGTTGGGTAACGCCAG | Reverse | — | pAM036 and PAM041 to amplify DNA template |

**Table S3. Transcription templates – scaffold components and PCR fragment sequences.**

| Name | Code | Type | Sequence from 5' to 3' | Label | Employment in Figure |
|---|---|---|---|---|---|
| Cy.5.5(-45/+45) | D021 | tDNA | TCGTTTCCAAACTCTTTTCGAACTTGTCTTCAACTGCTTTCGCATGAAGTACCTCCCAACTACTTTTCCTCACACTTGTACTCCA*T*G*A*C*T | Phosphoro-thioate (PTO, *) | Fig.1, Fig.S2, Fig.S3 |
| Cy.5.5(-45/+45) | D022 | ntDNA | [Cy5.5]AGTCATGGAGTACAAGTGTGAGGAAAAGTAGTTGGGAGGTACTTCATGCGAAAGCAGTTGAAGACAAGTTCGAAAAGAGTTTGGAAACGA | Cy5.5 | Fig.1, Fig.S2, Fig.S3 |
| (-45/+45)Cy5.5 | D030 | tDNA | [Cy5.5]TCGTTTCCAAACTCTTTTCGAACTTGTCTTCAACTGCTTTCGCATGAAGTACCTCCCAACTACTTTTCCTCACACTTGTACTCCATGACT | Cy5.5 | Fig.1, Fig.S2, Fig.S3(a) |
| (-45/+45)Cy5.5 | D031 | ntDNA | AGTCATGGAGTACAAGTGTGAGGAAAAGTAGTTGGGAGGTACTTCATGCGAAAGCAGTTGAAGACAAGTTCGAAAAGAGTTTGGA*A*A*C*G*A | PTO (*) | Fig.1, Fig.S2, Fig.S3(a) |
| (-45/+45)Cy5.5, -27G:C>T:A | D074 | tDNA | [Cy5.5]TCGTTTCCAAACTCTTTTCGAACTTGTCTTCAACTGCTTTCGCATGAAGTACCTCCCAACTACTTTTCCTCCCACTTGTACTCCATGACT | Cy5.5 | Fig.1, Fig.S3(a) |
| (-45/+45)Cy5.5, -27G:C>T:A | D075 | ntDNA | AGTCATGGAGTACAAGTGGGAGGAAAAGTAGTTGGGAGGTACTTCATGCGAAAGCAGTTGAAGACAAGTTCGAAAAGAGTTTGGA*A*A*C*G*A | PTO (*) | Fig.1, Fig.S3(a) |

| | | | | | |
|---|---|---|---|---|---|
| (-45/+45)Cy5.5, -28T:A>G:C | D076 | tDNA | [Cy5.5]TCGTTTCCAAACTCTTTTCGAACTTGTCTTCAACTGCTTTCGCATGAAGTACCTCCCAACTACTTTTCCTCAAACTTGTACTCCATGACT | Cy5.5 | Fig.1, Fig.S3(a) |
| (-45/+45)Cy5.5, -28T:A>G:C | D077 | ntDNA | AGTCATGGAGTACAAGTTTGAGGAAAAGTAGTTGGGAGGTACTTCATGCGAAAGCAGTTGAAGACAAGTTCGAAAAGAGTTTGGA*A*A*C*G*A | PTO (*) | Fig.1, Fig.S3(a) |
| (-80/-35)Cy5.5 | D054 | tDNA | [Cy5.5]ACTCCATGACTAAACCCCCCCTCCCATTACAAACTAAAATCTTACT | Cy5.5 | Fig.1, Fig.S3(a) |
| (-80/-35)Cy5.5 | D055 | ntDNA | AGTAAGATTTTAGTTTGTAATGGGAGGGGGGGTTTAGTCAT*G*G*A*G*T | PTO (*) | Fig.1, Fig.S3(a) |
| DNA(Cy3) | D094 | tDNA | CCCAACTACTTTTCCTCACACTTGTACTCCA | - | Fig.2a, Fig.3. Fig.S4(b-d), Fig.S5(a,b,d) |
| DNA(Cy3) | D095 | ntDNA | TGGAGTACAAGTGTGAGGAAAAGTAGTTGGG [Cy3] | Cy3 | Fig.2a, Fig.3, Fig.S4(b-d), Fig.S5(a-c) |
| DNA(Cy3, -28) | D097 | tDNA | CCCAACTACTTTTCCTCAAACTTGTACTCCA | - | Fig.3 (b-d), Fig.S5 (e) |
| DNA(Cy3, -28) | D098 | ntDNA | TGGAGTACAAGTTTGAGGAAAAGTAGTTGGG[Cy3] | Cy3 | Fig.3 (b-d), Fig.S5(e) |
| DNA(Cy3, -27) | D099 | tDNA | CCCAACTACTTTTCCTCCCACTTGTACTCCA | - | Fig.3 (b-d), Fig.S5(d) |
| DNA(Cy3, -27) | D100 | ntDNA | TGGAGTACAAGTGGGAGGAAAAGTAGTTGGG[Cy3] | Cy3 | Fig.3 (b-d), Fig.S5(d) |
| -90/+30 | D013 | tDNA | TTTCGAACTTGTCTTCAACTGCTTTCGCATGAAGTACCTCCCAACTACTTTTCCTCACACTTGTACTCCATGACTAAACCCCCCCTCCCATTACAAACTAAAATCTTACTTTTATTTTCT | - | Fig.4(a, b), Fig.6(b,d,e) |
| -90/+30 | D014 | ntDNA | AGAAAATAAAAGTAAGATTTTAGTTTGTAATGGGAGGGGGGGTTTAGTCATGGAGTACAAGTGTGAGGAAAAGTAGTTGGGAGGTACTTCATGCGAAAGCAGTTGAAGACAAGTTCGAAA | - | Fig.4(a, b), Fig.6(b,d,e) |
| -45/+45 | D082 | tDNA | TCGTTTCCAAACTCTTTTCGAACTTGTCTTCAACTGCTTTCGCATGAAGTACCTCCCAACTACTTTTCCTCACACTTGTACTCCATGACT | - | Fig.2(b,c),Fig.4 (a), Fig.5, Fig.S6 |
| -45/+45 | D083 | ntDNA | AGTCATGGAGTACAAGTGTGAGGAAAAGTAGTTGGGAGGTACTTCATGCGAAAGCAGTTGAAGACAAGTTCGAAAAGAGTTTGGAAACGA | - | Fig.2(b,c),Fig.4 (a), Fig.5, Fig.S6 |
| -30/+30 | D020 | tDNA | TTTCGAACTTGTCTTCAACTGCTTTCGCATGAAGTACCTCCCAACTACTTTTCCTCACAC | - | Fig.3(d), Fi.4(a,b,d), Fig.S4(b,d) |
| -30/+30 | D019 | ntDNA | GTGTGAGGAAAAGTAGTTGGGAGGTACTTCATGCGAAAGCAGTTGAAGACAAGTTCGAAA | - | Fig.3(d), Fi.4(a,b,d), Fig.S4(b,d) |
| -28/+30 | D056 | tDNA | TTTCGAACTTGTCTTCAACTGCTTTCGCATGAAGTACCTCCCAACTACTTTTCCTCAC | - | Fig.4(a,b,d), Fig.S4(d) |
| -28/+30 | D057 | ntDNA | GTGAGGAAAAGTAGTTGGGAGGTACTTCATGCGAAAGCAGTTGAAGACAAGTTCGAAA | - | Fig.4(a,b,d), Fig.S4(d) |
| -27/+30 | D058 | tDNA | TTTCGAACTTGTCTTCAACTGCTTTCGCATGAAGTACCTCCCAACTACTTTTCCTCA | - | Fig.4(a,b), |
| -27/+30 | D059 | ntDNA | TGAGGAAAAGTAGTTGGGAGGTACTTCATGCGAAAGCAGTTGAAGACAAGTTCGAAA | - | Fig.4(a,b), |
| -26/+30 | D049 | tDNA | TTTCGAACTTGTCTTCAACTGCTTTCGCATGAAGTACCTCCCAACTACTTTTCCTC | - | Fig.4(a,b), Fig.S4(d) |
| -26/+30 | D050 | ntDNA | GAGGAAAAGTAGTTGGGAGGTACTTCATGCGAAAGCAGTTGAAGACAAGTTCGAAA | - | Fig.4(a,b), Fig.S4(d) |
| -15/+30 | D018 | tDNA | TTTCGAACTTGTCTTCAACTGCTTTCGCATGAAGTACCTCCCAAC | - | Fig.4(a) |
| -15/+30 | D017 | ntDNA | GTTGGGAGGTACTTCATGCGAAAGCAGTTGAAGACAAGTTCGAAA | - | Fig.4(a) |
| -12/+30 | D024 | tDNA | TTTCGAACTTGTCTTCAACTGCTTTCGCATGTAAAGTACCTCCC | - | Fig.4(a) |

| | | | | | |
|---|---|---|---|---|---|
| -12/+30 | D025 | ntDNA | GGGAGGTACTTCATGCGAAAGCAGTTGAAGACAAGTTCGAAA | - | Fig.4(a) |
| -13del | D051 | tDNA | TTCCAAACTCTTTTCGAACTTGTCTTCAACTGCTTTCGCATGAAGTACCTCCCACTACTTTTCCTCACACTTGTACTCCATGACTAAACC | - | Fig.4(d), Fig.5(a,b), Fig.S4(d) |
| -13del | D052 | ntDNA | GGTTTAGTCATGGAGTACAAGTGTGAGGAAAAGTAGTGGGAGGTACTTCATGCGAAAGCAGTTGAAGACAAGTTCGAAAAGAGTTTGGAA | - | Fig.4(d), Fig.5(a,b), Fig.S4(d) |
| -13ins | D060 | tDNA | TTCCAAACTCTTTTCGAACTTGTCTTCAACTGCTTTCGCATGAAGTACCTCCCAAACTACTTTTCCTCACACTTGTACTCCATGACTAAACC | - | Fig.4(d), Fig.5(a,b), Fig.S4(d) |
| -13ins | D061 | ntDNA | GGTTTAGTCATGGAGTACAAGTGTGAGGAAAAGTAGTTTGGGAGGTACTTCATGCGAAAGCAGTTGAAGACAAGTTCGAAAAGAGTTTGGAA | - | Fig.4(d), Fig.5(a,b), Fig.S4(d) |
| -27T>G | D080 | tDNA | GAACTTGTCTTCAACTGCTTTCGCATGAAGTACCTCCCAACTACTTTTCCTCCCACTTGTACTCCATGACT | - | Fig.5 |
| -27T>G | D075 | ntDNA | — | PTO | Fig.5 |
| -28G>T | D081 | tDNA | TCGTTTCCAAACTCTTTTCGAACTTGTCTTCAACTGCTTTCGCATGAAGTACCTCCCAACTACTTTTCCTCAAACTTGTACTCCATGACT | - | Fig.5 |
| -28G>T | D077 | ntDNA | — | PTO | Fig.5 |
| FPt1 | D018 | tDNA | — | - | Fig.4(d), Fig.6(b,c,e), Fig.S4(d) |
| FPt1, FPt2 | D027 | ntDNA | TACTTCATGCGAAAGCAGTTGAAGACAAGTTCGAAA | - | Fig.4(d), Fig.6(b,c,e), Fig.S4(d) |
| FPt2 | D020 | tDNA | - | - | Fig.4(d), Fig.S4(d) |
| FPnt1, FPnt2 | D026 | tDNA | TTTCGAACTTGTCTTCAACTGCTTTCGCATGAAG | - | Fig.6(b,e) |
| FPnt1 | D025 | ntDNA | — | - | Fig.6(b,e) |
| FPnt2 | D017 | ntDNA | — | - | Fig.6(b,e) |
| Pb1 | D020 | tDNA | — | - | Fig.6(d,e) |
| Pb1 | D023 | ntDNA | GTGTGAGGAAAAGTAGTTGGCGTAGCAGGAGAAGTAAAGCAGTTGAAGACAAGTTCGAAA | - | Fig.6(d,e) |
| Pb2 | D08 | tDNA | TCGAATTCGTTTCCAAACTCTGTCTTCAACGGCTTTCTCATGAAGTACCTCCCAACTACTTTTCCTCACACTTGTACTCCATGACTAAACC | - | Fig.6(d,e,f) |
| Pb2 | D010 | ntDNA | GGTTTAGTCATGGAGTACAAGTGTGAGGAAAAGTAGTTGGGTCCATTCTATGGAGAAAGCCGTTGAAGACAGAGTTTGGAAACGAATTCGA | - | Fig.6(d,e,f) |
| -200/+150 | PCR product, and pAM036 | tDNA | TTAAGTTGGGTAACGCCAGGGTTTTCCCAGTCACGACGTTGTAAAACGACGGCCAGTGCCAAGCTTCACGCTGTATAGAGACTAGGCAGATCTGACGATCACCTAGCGACTCTCTCCACCGTTTGACGAGGCCATTTACAAAAACATAACGAACGACAAGCCTACTCGAATTCGTTTCCAAACTCTTTTCGAACTTGTCTTCAACTGCTTTCGCATGAAGTACCTCCCAACTACTTTTCCTCACACTTGTACTCCATGACTAAACCCCCCCCTCCCATTACAAACTAAAATCTTACTTTTATTTTCTTTTGCCCTCTCTGTCGCTCTGCCTTAACTACGTATTTCTCGCCGAGAAAAACTTCAATTTAAGCTATTCTCCAAAAATCTTAGCGTATATTTTTTTTCCAAAGTGACAGGGAGCTCGAATTCGTAATCATGTCATAGCTGTTTCCTGTGTGAAATTGTTATCCGCTCACAATTCCACACAACATACGAGCCG | - | Fig.6(g,h) |
| -200/+150 | PCR product, and pAM036 | ntDNA | CGGCTCGTATGTTGTGTGGAATTGTGAGCGGATAACAATTTCACACAGGAAACAGCTATGACATGATTACGAATTCGAGCTCCCTGTCACTTTGGAAAAAAAAATATACGCTAAGATTTTTGGAGAATAGCTTAAATTGAAGTTTTTCTCGGCGAGAAATACGTAGTTAAGGCAGAGCGACAGAGAGGGCAAAAGAAAATAAAAGTAAGATTTTAGTTTGTAATGGGAGGGGGGGTTTAGTCATGGAGTACAAGTGTGAGGAAAAGTAGTTGGGAGGTACTTCATGCGAAAGCAGTTGAAGACAAGTTCGAAAAGAGTTTGGAAACGAATTCGAGTAGGCTTGTCGTTCGTTATGTTTTTGTAAATGGCCTCGTC | - | Fig.6(g,h) |

| | | | | |
|---|---|---|---|---|
| | | | AAACGGTGGAGAGAGTCGCTAGGTGATCGTCAGATCTGCCT<br>AGTCTCTATACAGCGTGAAGCTTGGCACTGGCCGTCGTTTTA<br>CAACGTCGTGACTGGGAAAACCCTGGCGTTACCCAACTTAA | |
| non-specific DNA | PCR product, and pAM041 | tDNA | TTAAGTTGGGTAACGCCAGGGTTTTCCCAGTCACGACGTTGT<br>AAAACGACGGCCAGTGCCAAGCTTCTCCTATCAGGTATCTGT<br>CTTTGTTGTTGTGTCTGGACCTGGTGTTGTCTCTGTTGCTGTT<br>GCTGCTGCTGCTGCTGCTGTTGGGGCTGCTGCTGCTGTTGAA<br>ATTGTGGTTGGCGTTTCCCTATGTCTTGATCACTTTGTCCTTCT<br>TGTTCTGGCTGATTCTGGGGTCCTTGTGAGTTATGTGAGACA<br>GATGCATGTCGAGCTGAAAATAATTCGTCGTTTGTATTGTTA<br>ATTTCAAACTTCCAATCACGTCTGTATGATACCGAGACTCTTA<br>AGAGTCCAAACTTTGTCCATACAGGTGTGATCTTCTTTTGATC<br>TAAATGGGCTGGCAGGTTTGATTCGTTAAGCGCATTGGAATA<br>GAGCTCGAATTCGTAATCATGTCATAGCTGTTTCCTGTGTGA<br>AATTGTTATCCGCTCACAATTCCACACAACATACGAGCCG | - |
| non-specific DNA | PCR product, and pAM041 | ntDNA | CGGCTCGTATGTTGTGTGGAATTGTGAGCGGATAACAATTTC<br>ACACAGGAAACAGCTATGACATGATTACGAATTCGAGCTCTA<br>TTCCAATGCGCTTAACGAATCAAACCTGCCAGCCCATTTAGA<br>TCAAAAGAAGATCACACCTGTATGGACAAAGTTTGGACTCTT<br>AAGAGTCTCGGTATCATACAGACGTGATTGGAAGTTTGAAAT<br>TAACAATACAAACGACGAATTATTTTCAGCTCGACATGCATC<br>TGTCTCACATAACTCACAAGGACCCCAGAATCAGCCAGAACA<br>AGAAGGACAAAGTGATCAAGACATAGGGAAACGCCAACCAC<br>AATTTCAACAGCAGCAGCAGCCCCAACAGCAGCAGCAGCAG<br>CAGCAACAGCAACAGAGACAACACCAGGTCCAGACACAACA<br>ACAAAGACAGATACCTGATAGGAGAAGCTTGGCACTGGCCG<br>TCGTTTTACAACGTCGTGACTGGGAAAACCCTGGCGTTACCC<br>AACTTAA | - |

**Table S4. Exonuclease III footprinting summary**

| rDNA promoter region | Name of DNA oligonucleotides | Footprinting target | Footprinting direction | Footprint boundary, bp | Number of experiments |
|---|---|---|---|---|---|
| -45/+45 | Cy.5.5(-45/+45) | CF | downstream | -12 | 7 |
| -45/+45 | (-45/+45)Cy5.5 | CF | upstream | -32 (-33) | 7 |
| -45/+45, mutation at -27 bp | (-45/+45)Cy5.5, -27G:C>T:A | CF | upstream | none or -32 firmly supressed | 4 |
| -45/+45, mutation at -28bp | (-45/+45)Cy5.5, -28T:A>G:C | CF | upstream | -32 firmly supressed | 4 |
| -80/-35 | (-80/-35)Cy5.5 | CF | upstream nonspecific interaction | none | 4 |
| -45/+45 | Cy.5.5(-45/+45) | Pol1·Rrn3·CF complex | downstream | -12 | 3 |
| -45/+45 | Cy.5.5(-45/+45) | Pol1·Rrn3 complex | downstream | none | 3 |

**Table S5. Fit parameters.**

| Experiment type ID | Parameter | [CF] (nM) | DNA scaffolds | | | | Fit equations | Figure | Measurement method |
|---|---|---|---|---|---|---|---|---|---|
| | | | DNA(Cy3) fixed 24 nM | DNA(Cy3,-27) fixed 24 nM | DNA(Cy3,-28) fixed 24 nM | DNA(-45/45) titration 0–100 nM | | | |
| Measurements in FTB5 buffer | | | | | | | | | |
| 1 | $K_D^{app}$ (nM) | 12.5–200 | 106.3±63.6 | | | | 2 & 3 & 4 | 2a | Fluorometer |
| 1 | $F_{free}$ | | 35.6±0.9 | | | | 2 & 3 & 4 | 2a | Fluorometer |
| 1 | $F_{bound}$ | | 56.5±5.4 | | | | 2 & 3 & 4 | 2a | Fluorometer |
| 2 | $K_D^{app}$ (nM) | 50 (fixed) | | | | 20.5±4.5 | 2 & 3 & 4 | 2c | MP |
| 2 | $F_{free}$ | 50 | | | | 0 (fixed) | 2 & 3 & 4 | 2c | MP |
| 2 | $F_{bound}$ | 50 | | | | 1 (fixed) | 2 & 3 & 4 | 2c | MP |
| 3 | $K_D^{app}$ (nM) | 12.5–120 | 79±35 | 7.1±7.0 | 1.3±2.7 | | 2 & 3 & 4 | 3b | SF |
| 3 | $F_{free}$ | | 1 (fixed) | 1 (fixed) | 1 (fixed) | | 2 & 3 & 4 | 3b | SF |
| 3 | $F_{bound}$ | | 1.74±0.16 | 1.14±0.02 | 1.08±0.01 | | 2 & 3 & 4 | 3b | SF |
| 4 | $k_{obs}$ ($s^{-1}$) | 12.5<br>25<br>50<br>80<br>120<br><br>avg. | 0.0333±0.0008<br>0.0343±0.0009<br>0.0347±0.0005<br>0.0376±0.0004<br>0.0383±0.0003<br><br>0.036±0.002 | 0.133±0.008<br>0.112±0.005<br>0.138±0.003<br>0.152±0.004<br>0.180±0.003<br><br>0.143±0.023 | 0.661±0.143*<br>0.490±0.096*<br>0.533±0.053<br>0.312±0.034<br>0.281±0.020<br><br>0.375±0.112 | | 5 | 3a, 3c, S5c | SF |
| 4 | -*A* | 12.5<br>25<br>50<br>80<br>120 | 0.137±0.001<br>0.141±0.001<br>0.268±0.001<br>0.367±0.001<br>0.445±0.001 | 0.044±0.001<br>0.088±0.002<br>0.095±0.001<br>0.108±0.001<br>0.143±0.001 | 0.016±0.002<br>0.022±0.002<br>0.038±0.002<br>0.026±0.001<br>0.039±0.001 | | 5 | 3a, 3c, S5c | SF |
| 4 | *y0* | 12.5<br>25<br>50<br>80<br>120 | 1.130±0.001<br>1.134±0.001<br>1.258±0.001<br>1.364±0.001<br>1.432±0.001 | 1.045±0.001<br>1.097±0.002<br>1.095±0.001<br>1.105±0.001<br>1.143±0.001 | 1.016±0.001<br>1.022±0.001<br>1.038±0.001<br>1.026±0.001<br>1.039±0.001 | | 5 | 3a, 3c, S5c | SF |
| 4 | *m* (slope) | 12.5<br>25<br>50<br>80<br>120 | 0 (fixed)<br>0 (fixed)<br>0 (fixed)<br>0 (fixed)<br>0 (fixed) | $(1.17\pm0.17)\times10^{-4}$<br>$(1.33\pm0.25)\times10^{-4}$<br>$(1.34\pm0.15)\times10^{-4}$<br>$(1.73\pm0.18)\times10^{-4}$<br>$(2.68\pm0.15)\times10^{-4}$ | $(0.55\pm0.12)\times10^{-4}$<br>$(1.10\pm0.17)\times10^{-4}$<br>$(1.76\pm0.14)\times10^{-4}$<br>$(1.98\pm0.15)\times10^{-4}$<br>$(2.38\pm0.15)\times10^{-4}$ | | 5 | 3a, 3c, S5c | SF |
| 5 | *k* ($s^{-1}$) | | 0.0065±0.0001 | 0.0346±0.0001 | 0.2800±0.0033 | | 6 | 3d | SF |
| 5 | *A* | | 1.000±0.009 | 1.000±0.001 | 1.002±0.008 | | 6 | 3d | SF |
| 5 | *y0* | | $(-7.3\pm8000)\times10^{-6}$ | $(-3.1E\pm599)\times10^{-6}$ | -0.001± 0.002 | | 6 | 3d | SF |
| 5 | $\beta$ | | 0.713±0.004 | 0.916±0.003 | 0.779±0.011 | | 6 | 3d | SF |
| 5 | $k_{rev}$ ($s^{-1}$) | | 0.0076±0.0002 | 0.0358±0.0002 | 0.3107±0.0056 | | 8 | - | SF |
| 5 | $k_{for}$ ($s^{-1}$) | | 0.0284±0.0022 | 0.107±0.023 | 0.064±0.118 | | $k_{for}= k_{obs,avg}-k_{rev}$ | - | SF |
| 5 | $K_2$ | | 3.75±0.37 | 3.00±0.66 | 0.21±0.38 | | $K_2=k_{for}/k_{rev}$ | - | SF |
| Measurements in LTB5 buffer | | | | | | | | | |
| 6 | $K_D^{app}$ (nM) | 11.5–183.3 | 53.1±31.5 | | | | 2 & 3 & 4 | S4c | Fluorometer |
| 6 | $F_{free}$ | | 32.9±31.5 | | | | 2 & 3 & 4 | S4c | Fluorometer |
| 6 | $F_{bound}$ | | 57.7±4.5 | | | | 2 & 3 & 4 | S4c | Fluorometer |

Experiment IDs
1: Equilibrium binding of CF to Cy3-labelled rDNA promoters, binding measured by fluorometer.
2: Equilibrium binding of CF to non-labelled rDNA promoter, binding measure with mass photometer.
3: Equilibrium binding of CF to Cy3-labelled rDNA promoters, binding measured by stopped flow.
4: Kinetic rate of CF binding to Cy3-labelled rDNA promoters, binding measured by stopped flow.
5: Kinetic rate of CF dissociation from Cy3-labelled rDNA promoters, binding measured by stopped flow.
6: Equilibrium binding of CF to WT Cy3-labelled rDNA promoter in LTB5 buffer, binding measured by fluorometer.

nb. experiment types 1–5 were done in FTB5 buffer.

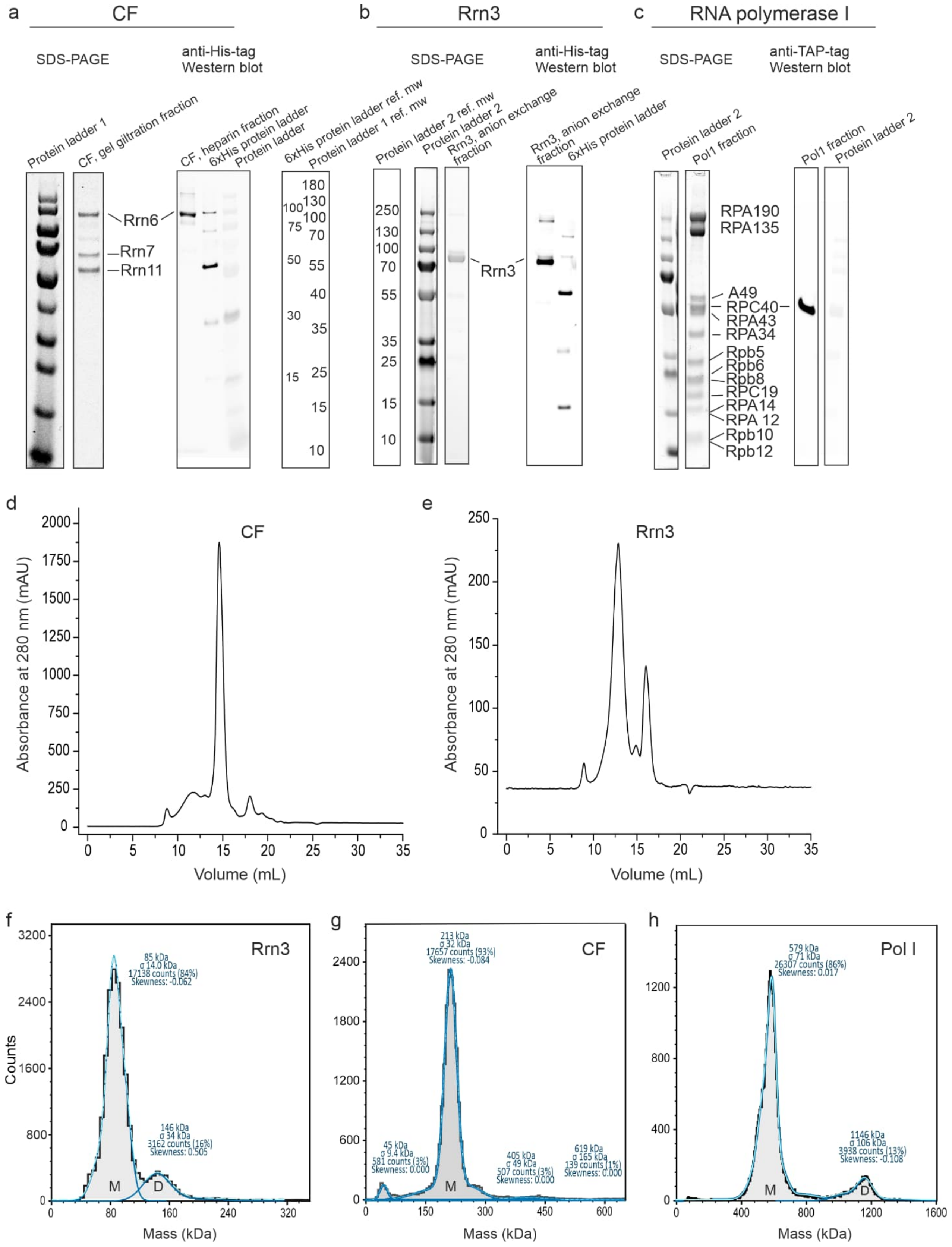

**Supplementary Figure 1. Characterization of purified RNA polymerase I and transcription factors.** Analysis of purified (a) CF, (b), Rrn3 and (c) RNA polymerase I proteins using SDS-PAGE and Western analysis. Protein ladder 1 (PageRuler™ prestained protein ladder, ThermoFisher Scientific), protein ladder 2(PageRuler™ Plus prestained protein ladder, ThermoFisher Scientific) and 6xHis protein ladder (6xHis Protein Ladder, Qiagen) were used as molecular weight standards. Size-exclusion purification chromatograms of (d) CF and (e) Rrn3. Apparent molecular weight (mean±σ; skewness) and molecular distribution (counts of molecules, %) obtained from representative mass photometer histograms of (f) Rrn3, (g) CF and (h) Pol I are indicated in blue font. The inferred protein monomer and protein dimer populations are marked with M and D, respectively. Three independent measurements were performed for each protein.

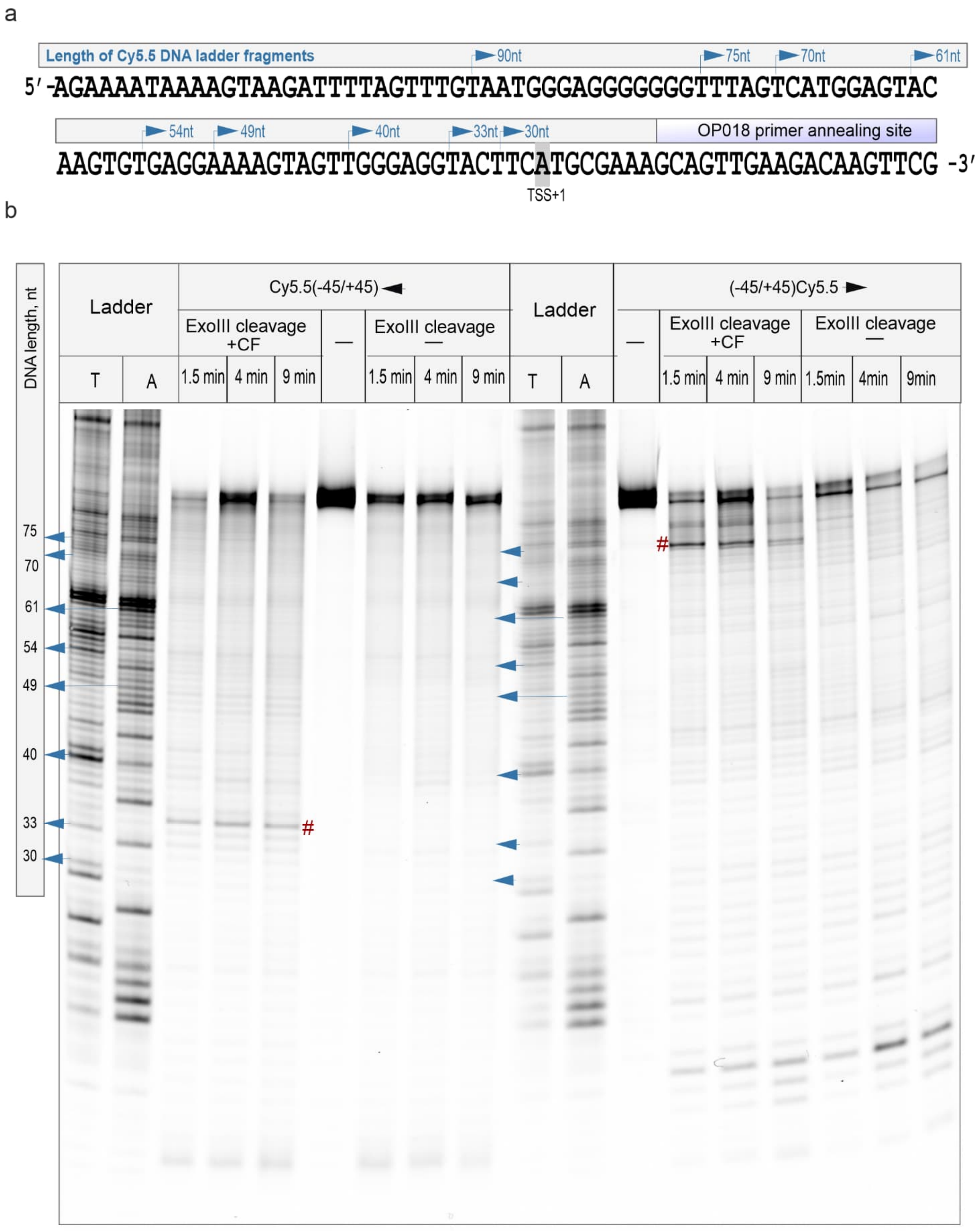

**Supplementary Figure 2. Calibration and time-dependence of exonuclease III footprinting.** (a) The schematic representation of the control DNA scaffold used as a template to produce DNA calibration ladder for exoIII footprinting and transcription start site experiments. The scaffold contains -90/+30 fragment of rDNA promoter; the sequence of non-template DNA strand from -90 to +27 is shown. The ladder was produced by Sanger sequencing reaction using a 19 nucleotides long primer (OP018), which binds to the +9/+27 region of the scaffold and which is labelled with cyanine 5.5 (Cy5.5) fluorophore at the 5' end. Blue arrows on the scaffold sequence and gel scan indicate the 3' ends and electrophoretic mobility of specific length DNA ladder strands. (b) Time-dependence of exoIII driven cleavage of rDNA promoter (span -45/+45) in the presence (+CF) and absence (-) of CF is shown. Black left and right arrows indicate the experiments to map either the downstream or upstream edge of the CF footprint, respectively. CF footprint is highlighted with # symbol.

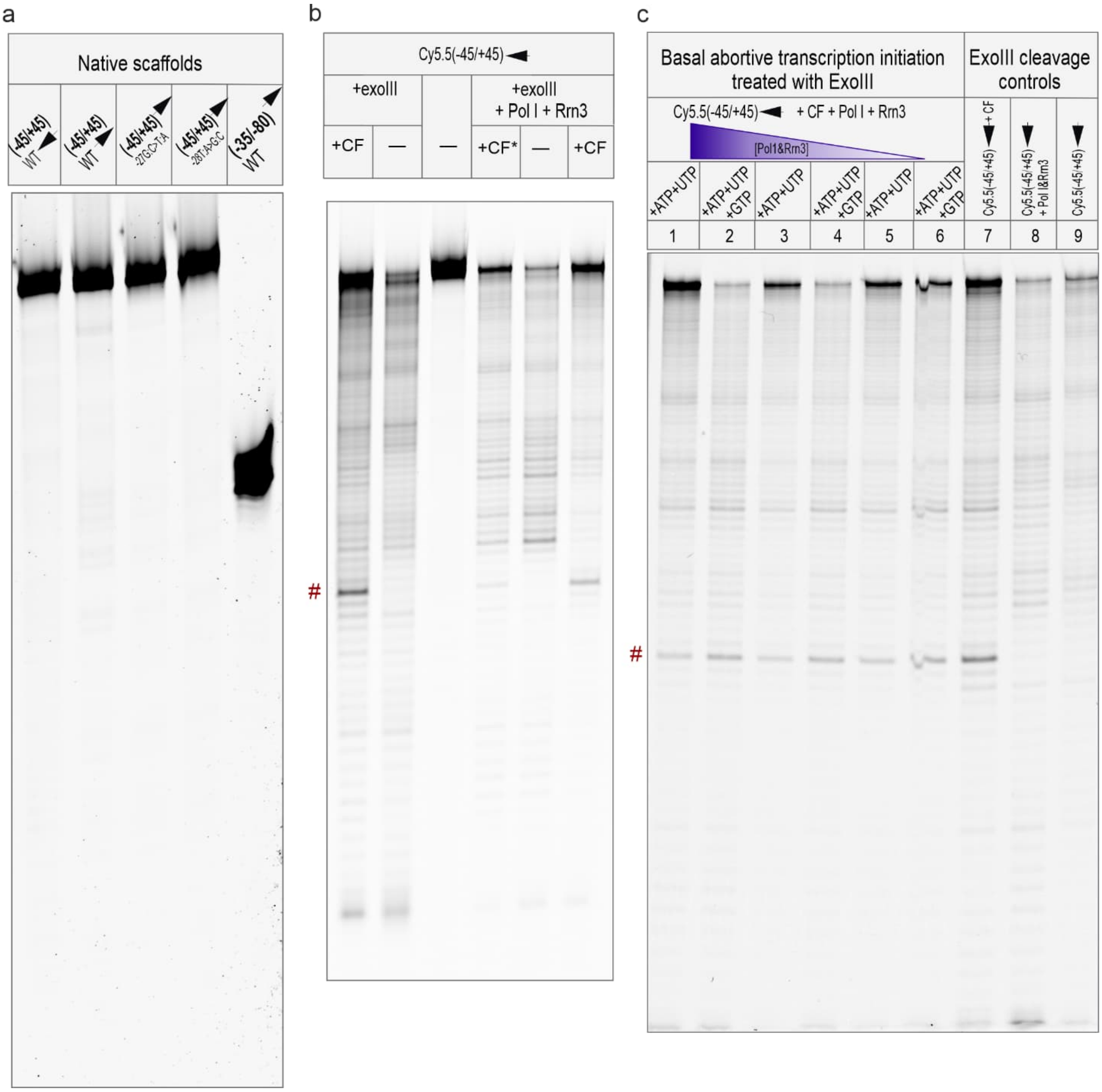

**Supplementary Figure 3. Exonuclease III footprinting of basal transcription initiation complex.** (a) PAGE analysis of untreated rDNA promoter templates used in exoIII footprinting experiments demonstrate their excellent integrity and purity. Scaffolds were run on the same gel with data presented in Figure 1. Left and right arrow indicate the promoters used for footprinting from the downstream and upstream direction, respectively. The span, e.g., (-45/+45), and relevant base substitutions in the promoters are shown. (b) The basal transcription preinitiation complex was footprinted from the downstream direction on rDNA promoter (-45/+45). The assembly samples consisted of 10 nM DNA-100 nM CF (lane 1); cleavage and integrity controls: 10 nM DNA treated with exoIII (lane 2) and 10 nM DNA alone (lane 3); 10nM DNA-50nMCF-100nM Pol1-400 nM Rrn3 (lane 4); 10nM DNA-100nM-400 nM Rrn3 (lane 5) and 10 nM DNA-100 nM CF-100 nM Pol1-400 nM Rrn3 (lane 6). (c) The basal transcription initiation complex during abortive initiation was exoIII footprinted from the downstream direction. The assembled complexes consisted of 10 nM rDNA promoter (-45/+45), 150 nM CF, and 20–50 nM holoenzyme (1:4 Pol I:Rrn3 mixture). The first set of samples containing 50 nM (sample lane 1), 30 nM (lane 3) or 20nM (lane 5) holoenzyme, respectively, were activated with 500 nM ATP and UTP nucleotides in ratio 1:1. The second set of samples containing 50 nM (lane 2), 30 nM (lane 4) or 20nM (lane 6) holoenzyme, respectively, were activated with 500 nM ATP, UTP and GTP in ratio 1:1. Cleavage controls included the promoter supplemented with 150 nM CF alone (lane 7), 30 nM holoenzyme alone (lane 8) or none (lane 9).

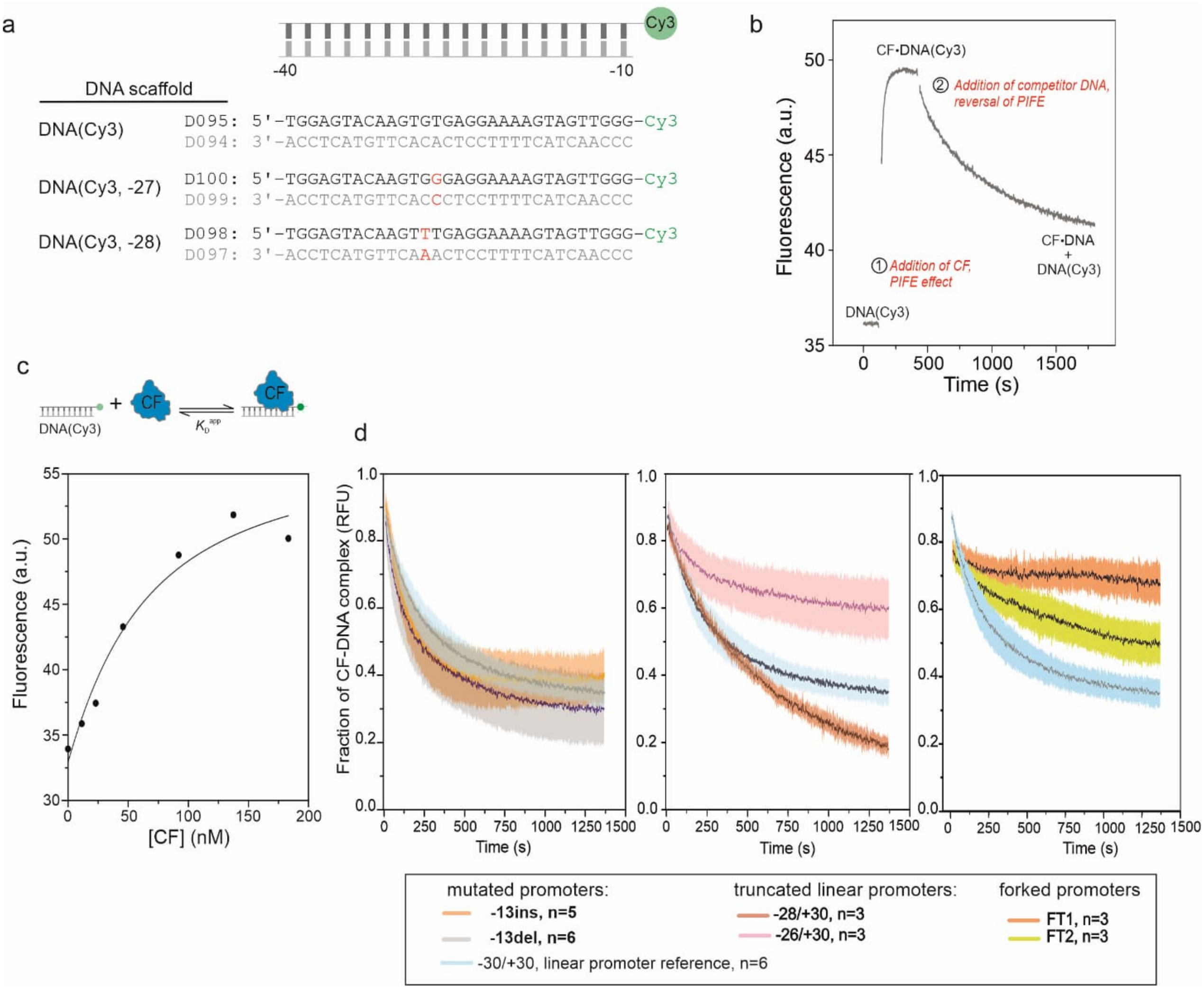

**Supplementary Figure 4**. **Binding and dissociation of CF from rDNA promoter using spectrofluorometer detection.** (a) The schematic structure and sequences of promoter DNAs used in PIFE assays are shown. Non-template and template DNA strands are indicated in dark and light grey, respectively. The mutated base pairs at positions -27 or -28 are in red font. Cy3-fluorophore is linked to the 3' end of non-template DNA. Dxxx numbers refer to the systematic names of DNA oligo in our laboratory. (b) Fluorescence intensity of 24 nM Cy3-labelled rDNA promoter [DNA(Cy3)] was monitored in spectrofluorometer. The addition of 120 nM CF to the reaction cuvette leads to the formation of CF·DNA(Cy3) complex and increase in fluorescence intensity (marked with 1). Further addition of 227 nM unlabeled (-30/+30) DNA promoter triggers CF dissociation from DNA(Cy3) and decrease in fluorescence intensity (marked with 2). (c) Apparent dissociation constant of CF·DNA(Cy3) complex was determined in lowered ionic strength buffer LTB5. The theoretical curve was obtained by fitting Equations 2–4 to data (n=1). (d) Normalized dissociation traces of preformed CF·DNA(Cy3) (the trace is average of n experiments and its SE is shown as transparent cloud) in PIFE competition assay. Increment of fluorescence intensity between DNA(Cy3) alone and CF-DNA(Cy3) mixture was set to 1 in each record. Dissociation fluorescence intensities were then converted to values within the range 0–1. The dissociation reaction was initiated by the addition of control native promoter (-30/+30), the promoters carrying single nucleotide deletion or insertion at position -13 (left panel), the promoters with upstream truncated at -28 or -26 (middle panel), or the fork promoters FPt1 or FPt2 (right panel). Data was recorded using a spectrofluorometer.

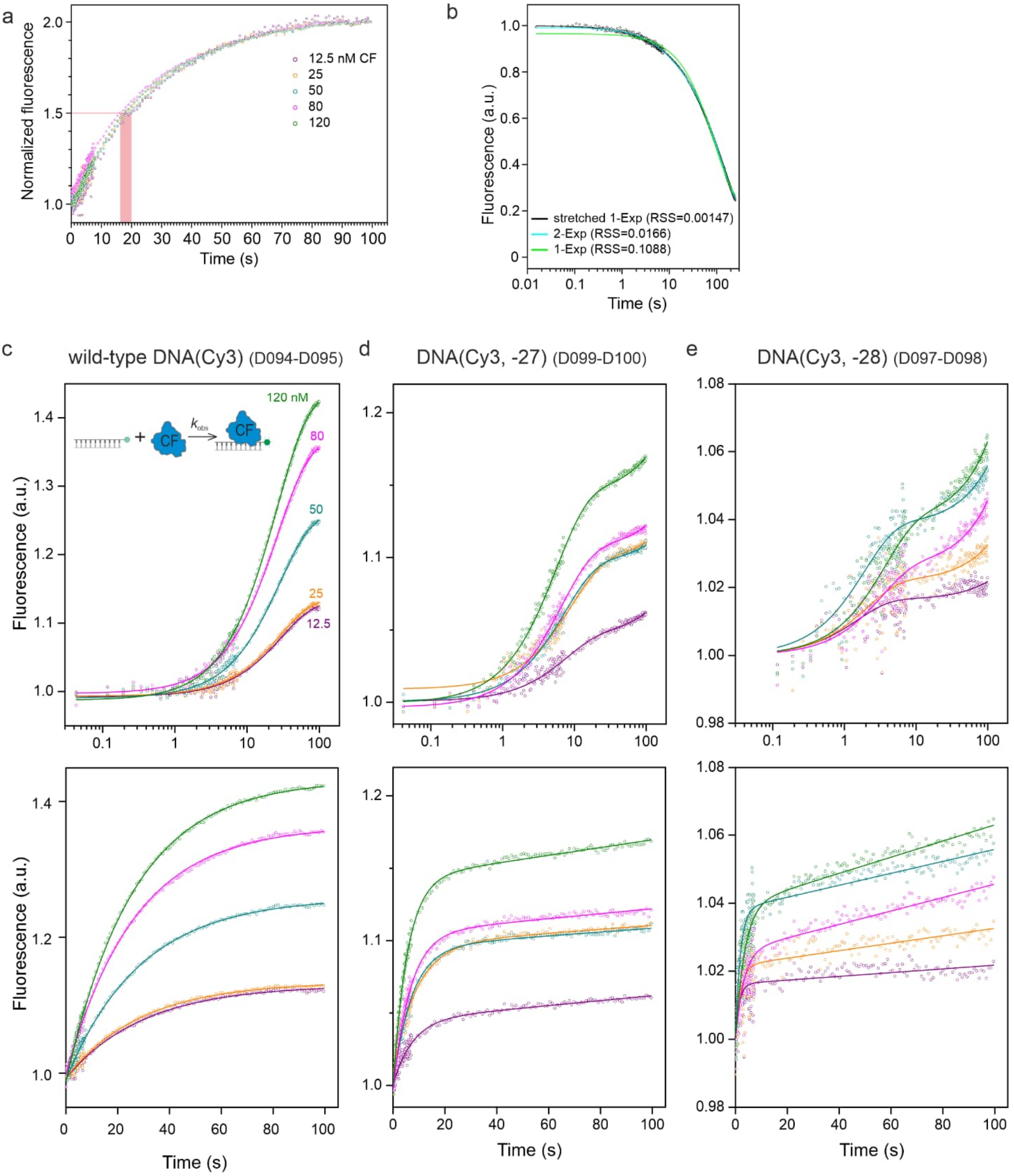

**Supplementary Figure 5. Binding and dissociation of CF from rDNA promoter in stopped flow experiments.** (a) Stopped flow traces monitoring the binding of CF to DNA(Cy3) were normalized to equal amplitudes. Pink shading highlights approximated half-lives (about 16–20 s) of the reactions. (b) SF trace monitoring the dissociation of CF from WT DNA(Cy3) was fit using single exponential, double exponential or stretched single exponential (Eq. 6) equation, respectively. Small RSS (residual-sum-of-squares) indicates better fit. Fluorescence intensity trajectories report the binding of CF (used as 12.5–120 nM) to (c) wild-type DNA(Cy3) [same data is shown as normalized in panel a], (d) DNA(Cy3, -27), or (e) DNA (Cy3, -28) promoter scaffolds. Data is shown in both logarithmic (top) and linear (bottom) time scale. DNA concentration was 24 nM in each experiment. The fit curves were obtained using Eq. 5 and parameter values in Table S5.

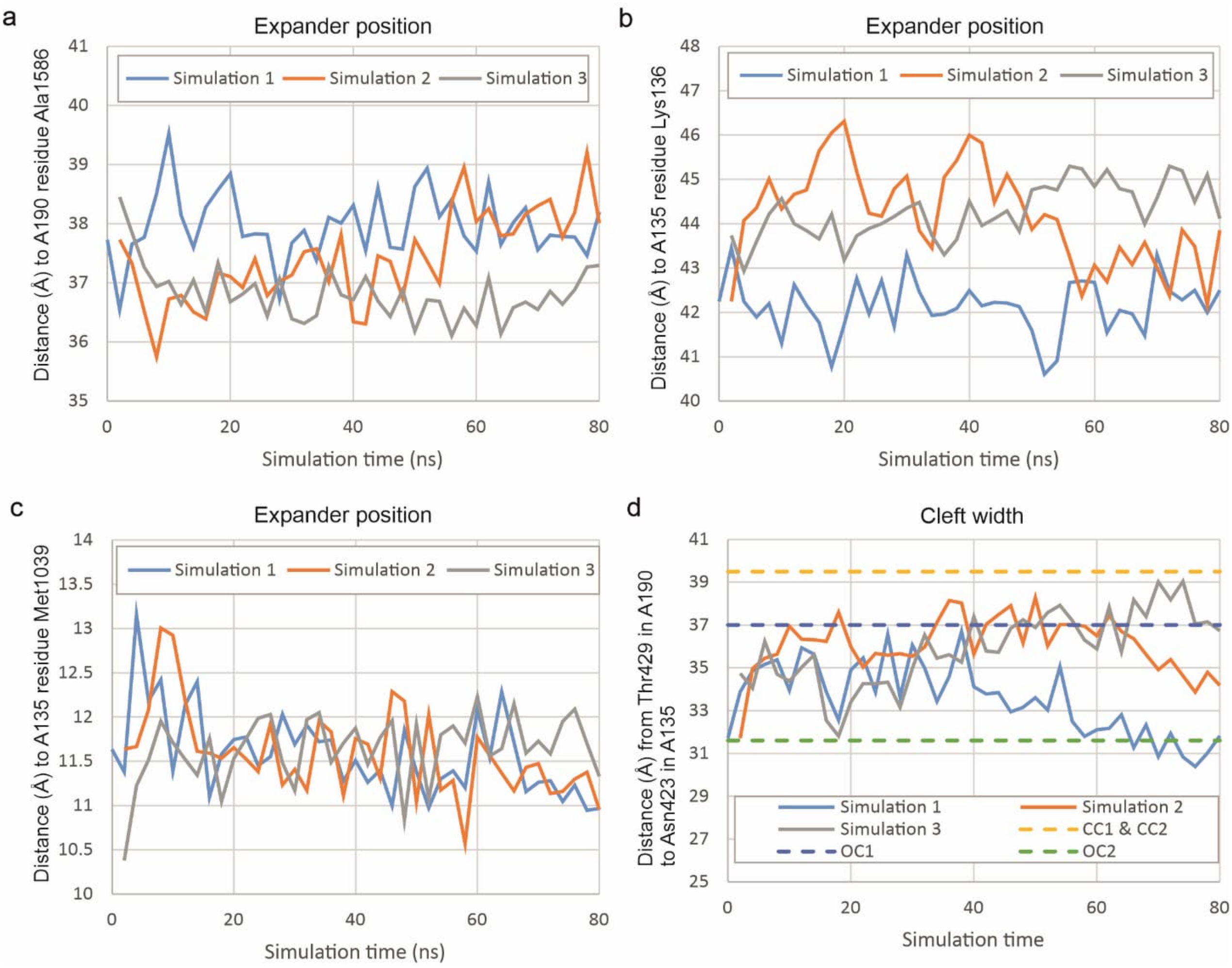

**Supplementary Figure 6. Analysis of Pol I expander position and cleft width in MD trajectories.** The stable docking of expander in the DNA-binding cleft of Pol I is demonstrated by measuring the distance from A190 residue Ser1374 (locates to the middle of expander helix) to three directions represented by residues (a) Ala1586 in A190, (b) Lys136 in A135, and (c) Met1039 in A135. Cα atoms were used in the distance measurements. (d) The width of Pol I cleft was estimated by measuring the distance between Cα atoms of Thr429 in A190 and Asn423 in A135. For comparison, also the distances from the cryo-EM based models of Pol I closed and open complexes are shown. Wild-type promoter was used in the simulations.

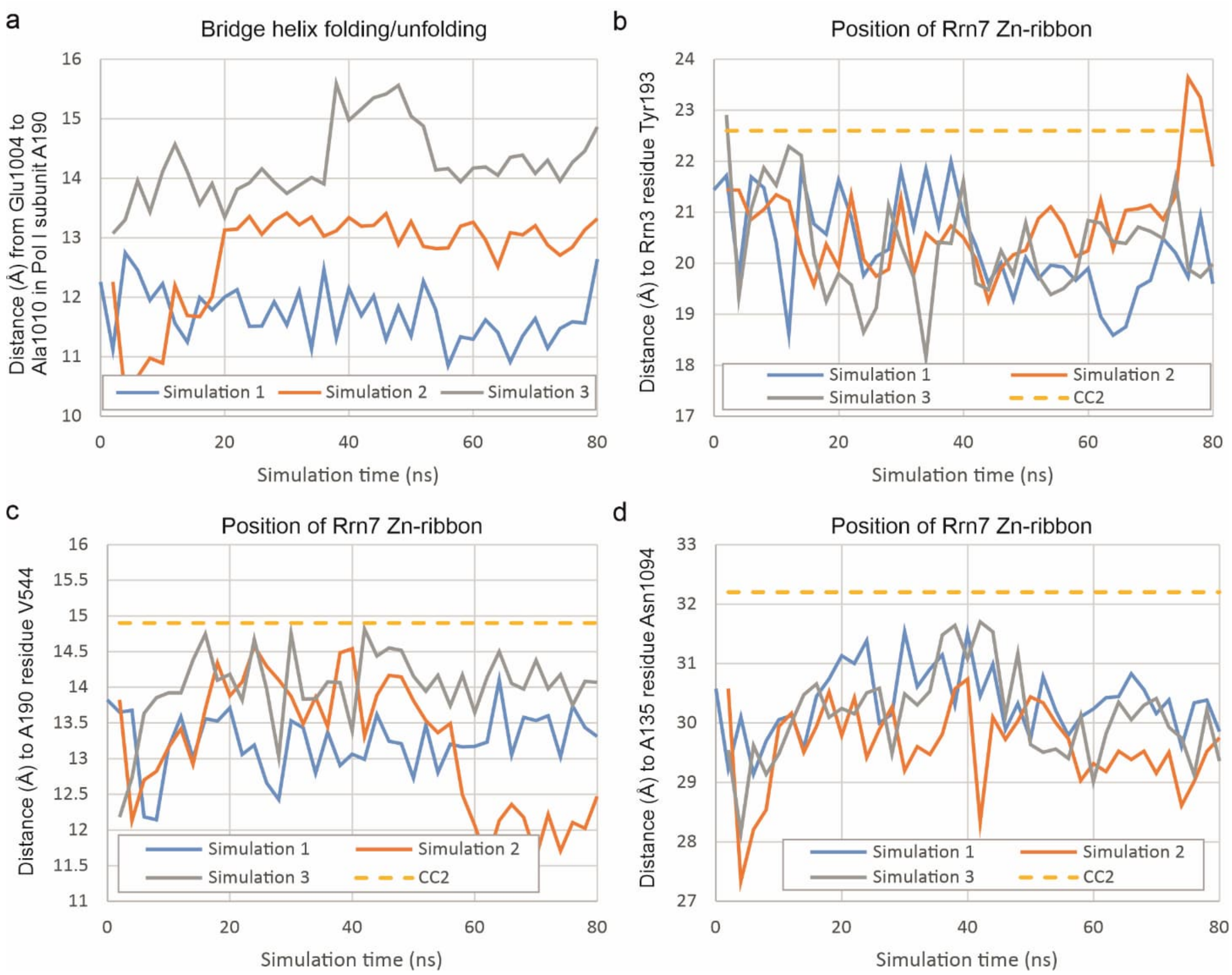

**Supplementary Figure 7. Analysis of Pol I bridge helix folding dynamics and Rrn7 Zn-ribbon position in MD trajectories.** (a) The folding and unfolding of bridge helix was demonstrated by the decrease and increase in the distance between residues Glu1004 and Ala1010 in A190 subunit of Pol I, respectively. The stable position of Zn-ribbon domain of Rrn7 (CF subunit) was inferred by measuring the distance from its Ser17 to three directions represented by residues (a) Tyr193 in Rrn3, (b) Val544 in A190, and (c) Asn1094 in A135. Cα atoms were used in the distance measurements. Wild-type promoter was used in the simulations.

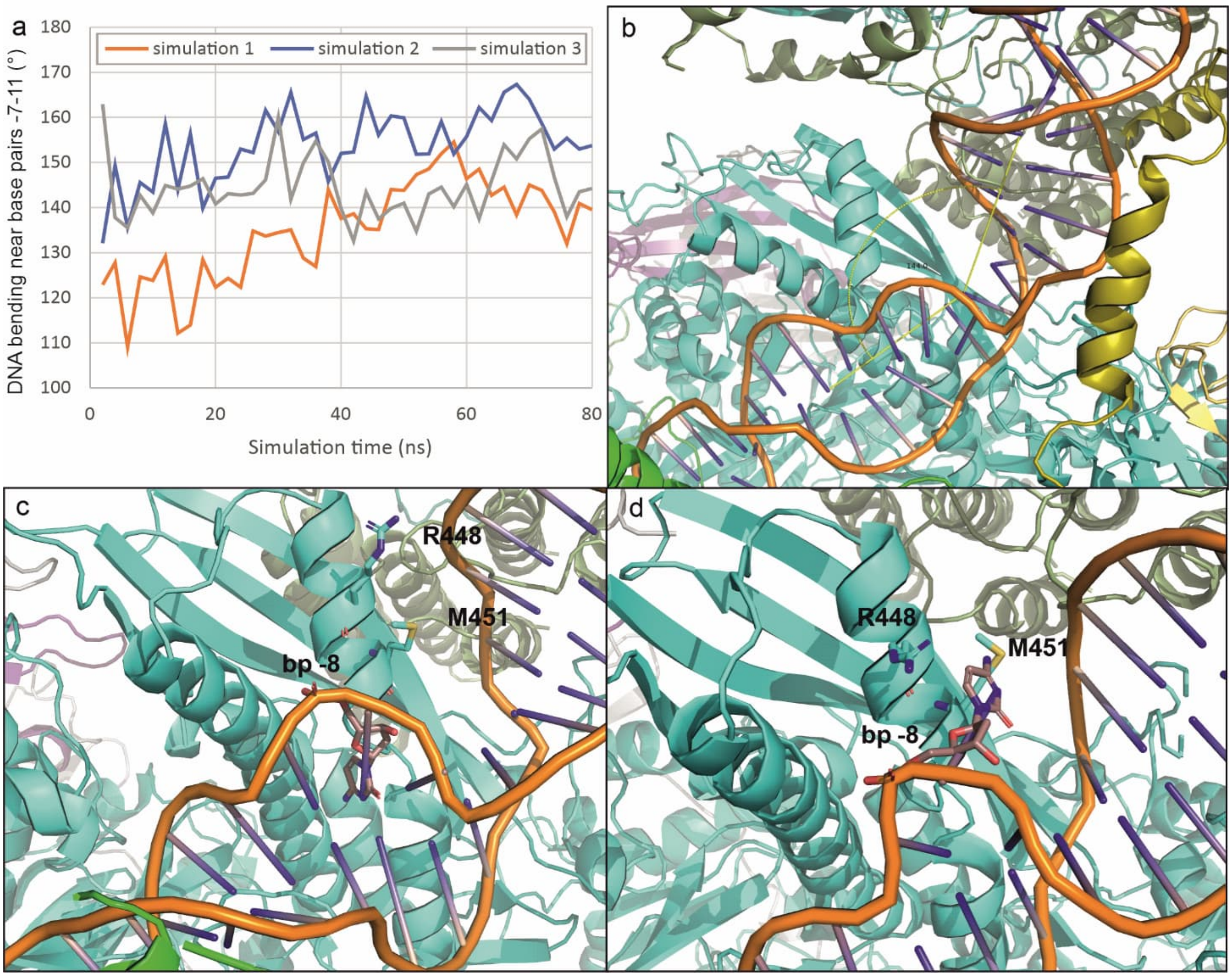

**Supplementary Figure 8. Kinking and unstable base pairing in the rDNA promoter region from -11 to -6 in MD trajectories.** (a) The kinking of rDNA promoter near positions -11 and -7 in three MD simulations is shown. (b) Trajectory snapshot demonstrates how the promoter kinking was measured by determining the angle between promoter positions -6, -10 and -15. (c) The snapshot demonstrates canonical base pairing like conformation of bp -8 in simulation 3. (d) Base pair -8 in fully flipped out conformation, interacting with residues Arg448 and Met451 of A135 protrusion domain in simulation 2. Wild-type promoter was used in the simulations.

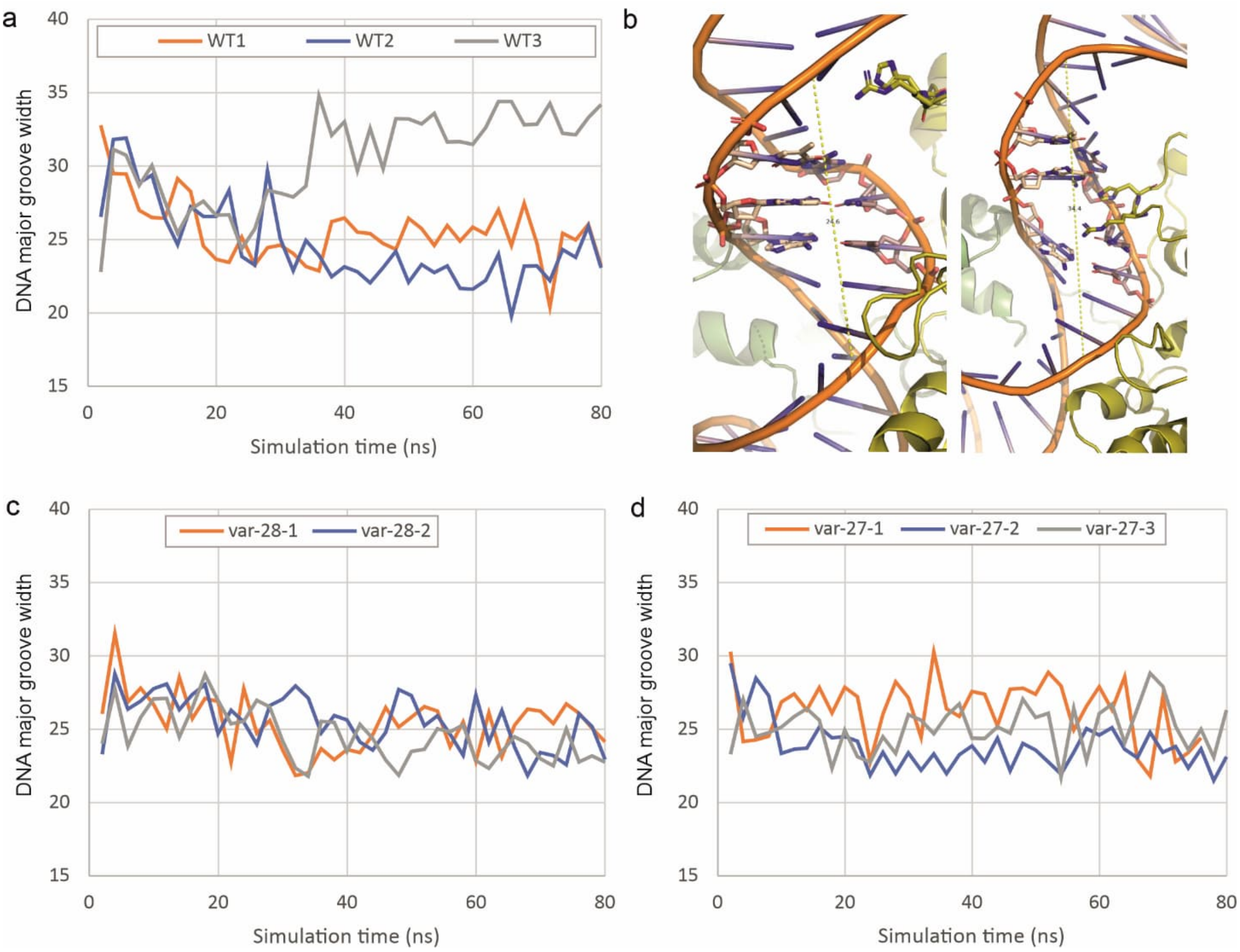

**Supplementary Figure 9. The width of rDNA promoter major groove from -27 to -22 and the insertion of the loop between α7 and α8 of Rrn7 in MD trajectories.** (a) The DNA major groove width in different MD simulations using wild-type promoter. (b) The groove width was estimated by measuring the distance between the phosphate atoms of bp -28 (non-template strand) and -21 (template strand) with example frames shown from WT simulation 1 (left, narrow groove width of 24.6 Å) and WT simulation 3 (right, wide groove of 34.4 Å). The residues Arg293 and His294 in the loop between α7 and α8 of Rrn7 (CF) are shown as stick models and coloured with yellow and blue. The DNA major groove width in simulations performed using the promoter mutated (c) at bp -28 or (d) at bp -27.
